# Physiology-forward identification of bile acid sensitive vomeronasal receptors

**DOI:** 10.1101/766592

**Authors:** Wen Mai Wong, Jie Cao, Xingjian Zhang, Wayne I. Doyle, Luis L. Mercado, Laurent Gautron, Julian P. Meeks

## Abstract

The mouse accessory olfactory system (AOS) supports social and reproductive behavior through the sensation of environmental chemosignals. A growing number of excreted steroids have been shown to be potent AOS cues, including bile acids (BAs) found in feces. As is still the case with most AOS ligands, the specific receptors used by vomeronasal sensory neurons (VSNs) to detect BAs remain unknown. To identify VSN BA receptors, we first performed a deep analysis of VSN BA tuning using volumetric GCaMP6f/s Ca^2+^ imaging. These experiments revealed both broadly and narrowly tuned populations of BA-receptive VSNs with sub-micromolar sensitivities. We then developed a new physiology-forward approach for identifying AOS ligand-receptor interactions, which we call <u>F</u>luorescence <u>L</u>ive Imaging for <u>C</u>ell <u>C</u>apture and <u>R</u>NA<u>-seq</u>, or FLICCR-seq. FLICCR-seq analysis revealed 5 specific V1R-family receptors enriched in BA-sensitive VSNs. These studies introduce a powerful new approach for ligand-receptor matching and reveal biological mechanisms underlying mammalian BA chemosensation.

## Introduction

Chemosensory systems extract salient information from environmental cues that are critical for social and reproductive behaviors. In mice and other terrestrial mammals, the accessory olfactory system (AOS) guides innate behaviors such as territorial aggression and mating (Dulac and Torello, 2003; Keller et al., 2009). The initial detection of olfactory stimuli in the AOS is mediated by vomeronasal sensory neurons (VSNs), located in the vomeronasal organ (VNO), before these signals are routed to the accessory olfactory bulb and then behaviorally important regions including the medial amygdala and bed nucleus of the *stria terminalis*.

Though the behavioral impacts of AOS chemosensation are manifold, our knowledge of the full complement of natural ligands for VSNs remains incomplete. Natural AOS ligands are found in the excretions of conspecific and heterospecific animals (*e.g*., urine, feces, tears, and saliva). These natural ligands include, but are not limited to, polar steroids, bile acids, major urinary proteins, and major histocompatibility complex peptide ligands (Chamero et al., 2007; Doyle et al., 2016; Leinders-Zufall et al., 2004; Nodari et al., 2008). As we learn more about the full natural complement of AOS ligands, new questions are arising about how these cues are detected by specific types of VSNs. Such knowledge is critical as we seek to understand how patterns of VSN activation by complex blends of environmental chemosignals cause changes in animal physiology and behavior. A major barrier preventing a deeper understanding of AOS function is a lack of ligand-receptor matches for AOS cues. VSNs generally express just one or two of ∼300 unique vomeronasal receptors (VRs) or formyl peptide receptors (Dulac and Torello, 2003; Martini et al., 2001). The largely monoallelic expression of individual VRs by VSNs means that each VSN takes on the chemosensory sensitivity (or “tuning”) of the dominantly expressed receptor. In theory, this feature would greatly simplify the process of unambiguously matching VSN ligands to their cognate receptors. However, efforts to generate large-scale VR screening assays via heterologous expression has, for the most part, been unsuccessful (though see (Dey et al., 2013)). Other efforts have explored VNO ligand-receptor interactions using various approaches (Haga-Yamanaka et al., 2015; Isogai et al., 2011; Lee et al., 2019; Stein et al., 2016). Despite recent progress, a small fraction of the VR family has confirmed ligands (Haga-Yamanaka et al., 2014; Haga et al., 2010; Isogai et al., 2011; Isogai et al., 2018; Lee et al., 2019; Osakada et al., 2018).

Given the importance of AOS ligand-receptor interactions for mammalian physiology and social behavior, we sought to develop a pipeline for matching AOS ligands to receptors. We devised a physiology-forward approach using native VNO tissue (with endogenous VR expression) to identify the receptors for 4 recently-discovered natural bile acid (BA) ligands present in feces (Doyle et al., 2016). BAs are steroids that are structurally similar to other potent AOS ligands including sulfated and carboxylated glucocorticoids (Fu et al., 2015; Nodari et al., 2008). Despite evidence that many VSNs show selectivity for BAs compared to these other steroids, and evidence that some VRs are exquisitely sensitive to other steroids (Fu et al., 2015; Haga-Yamanaka et al., 2015; Isogai et al., 2011; Lee et al., 2019), questions remain about whether there are BA specialist VRs or whether BAs nonselectively activate certain steroid-sensitive VRs.

Here, we report the discovery of distinct populations of VSNs that are BA specialists and others that are BA generalists. We comprehensively mapped the sensory space of VSNs to four prominent BAs and several sulfated steroids, including sex steroids and glucocorticoids, finding that some BA generalists are also sensitive to sulfated glucocorticoids. We report the successful development of a function-forward strategy, which we term <u>F</u>luorescence <u>L</u>ive Imaging and <u>C</u>ell <u>C</u>apture for <u>R</u>NA-<u>seq</u> (FLICCR-seq), to isolate highly BA-sensitive VSNs for single-cell RNA sequencing. We used FLICCR-seq show that BA-sensitive VSNs are enriched in the expression of at least five V1R-family VRs, two of which appear to be associated with broad BA tuning, and three that are associated with selective detection of specific BAs. Collectively, these studies reveal mechanisms of mammalian BA chemosensation and improve our understanding of chemosensory information as it enters the behaviorally-important AOS pathway.

## Methods

### Animals

All animal procedures were performed in accordance with the Institutional Animal Care and Use Committee at the University of Texas Southwestern Medical Center and follow guidelines from the National Institutes of Health. Physiological experiments were performed with *OMP*^*tm4(cre)Mom*^*/*J knock-in mice (*OMP-Cre* mice; Jackson Laboratory Stock #006668) mated to either *Gt(ROSA)26Sor*^*tm96(CAG-GCaMP6s)Hze*^/J mice (Ai96 mice; Jackson Laboratory Stock #024106) or CG-*Igs7*^*tm148.1(tet0-GCaMP6f,CAG-tTA2)Hze*^*/J* mice (Ai148 mice; Jackson Laboratory Stock #030328). These mice express the genetically encoded Ca^2+^ indicator GCaMP6s (OMP-Cre^+/-^, Ai96^+/-^) or GCaMP6f (OMP-Cre^+/-^, Ai148^+/-^) in VSNs, further referred to as OMPxAi96 mice or OMPxAi148 mice, respectively. Mice were between 6 and 15 weeks of age. The number, strain, and sex of the animals used in each experiment are described in corresponding figure legends. BaseScope experiments were performed with wild-type C57Bl/6J animals.

### Solutions and stimulus presentation

All reagents were purchased from Sigma-Aldrich (St. Louis, MO, USA) unless otherwise specified. Bile acid stimuli included cholic acid (CA), deoxycholic acid (DCA), chenodeoxycholic acid (CDCA) and lithocholic acid (LCA). Sulfated steroids included epitestosterone-17 sulfate (A6940), 17α-estradiol-3-sulphate (E0893), 5α-pregnan-3β-ol-20-one sulfate, (P3865) and corticosterone-21-sulfate (Q1570) purchased from Steraloids, Inc. (Newport, RI, USA). Stock solutions (20 mM) of all BAs and sulfated steroids were prepared in methanol and diluted to their final concentration in Ringer’s solution containing (in mM): 115 NaCl, 5 KCl, 2 CaCl_2_, 2 MgCl_2_, 25 NaHCO_3_, 10 HEPES and 10 glucose. Artificial cerebrospinal fluid (aCSF) contained (in mM): 125 NaCl, 2.5 KCl, 2 CaCl_2_, 1 MgCl_2_, 25 NaHCO_3_, 1.25 NaH_2_PO_4_, 25 glucose, 3 myo-inositol, 2 sodium pyruvate and 0.5 sodium ascorbate. All sulfated steroids were diluted to 10 µM (1:2000) for experiments. BAs were diluted to a range of concentrations from 0.1 µM to 10 µM. Control stimuli consisted of Ringer’s solution with 1:2000 methanol (the highest concentration of methanol in any individual stimulus). Stimuli were applied for 15 s using an air pressure-driven reservoir via a 16-in-1 multi-barrel ‘perfusion pencil’ (Automate Scientific, Berkeley, CA, USA).

### Volumetric VNO Ca^2+^ imaging

Following deep isofluorane anesthesia and rapid decapitation, VNOs were dissected and the vomeronasal epithelium carefully removed under a dissection microscope (Leica Microsystems, Buffalo Grove, IL, USA) similar to previous studies (Turaga and Holy, 2012; Wong et al., 2018). The vomeronasal epithelium was mounted onto nitrocellulose paper (Fisher Scientific, Atlanta, GA, USA) before placement into a custom imaging chamber. Volumetric Ca^2+^ imaging was performed using a custom objective-coupled planar illumination (OCPI) microscope (Holekamp et al., 2008) with refinements described previously (Wong et al., 2018). GCaMP6s/f fluorescence was acquired using custom software that synchronized imaging with stimulus delivery via a randomized, interleaved stimulus delivery system (Automate Scientific). Images stacks containing 51 frames and spanning ∼700 µm laterally, 250-400 µm axially, and ∼150 µm in depth were taken once every 3 s (∼ 0.33 Hz). Each individual stimulus was delivered for 5 consecutive stacks (∼15 seconds) with at least 10 stacks between stimulus trials (≥30 seconds). Each analyzed experiment completed at least three complete randomized, interleaved stimulus blocks. Stimulation patterns are further described in the Results.

### Data analysis of volumetric VNO Ca^2+^ imaging

Data analysis was performed using custom MATLAB software similar to previous studies (Hammen et al., 2014; Turaga and Holy, 2012). Briefly, image stacks underwent a rigid registration followed by a nonrigid warping. Then ΔF/F, the relative change in GCaMP6s/f intensity, was calculated by subtracting the mean voxel intensity in three consecutive pre-stimulus stacks from the mean voxel intensity of three stacks during stimulus delivery, then dividing by the value of the mean pre-stimulus intensity. Volumetric ROIs were manually drawn around the cell bodies of well-registered VSNs that reliably responded to stimulation as well as spontaneously active neurons. Following the volumetric ROI selection, the mean voxel intensity for each ROI was calculated for every image stack in the experiment (∼1,200 stacks) generating a matrix of fluorescence intensity. The across-trial mean ΔF/F for all ROIs and stimulus applications was calculated using this matrix. The ΔF/F of all ROIs in response to stimuli were then clustered using a bootstrapping clustering method based on the mean shift method (Comaniciu and Meer, 2002)(Meeks et al., 2010). This identifies common patterns of activity in the population and group them together.

### *Ex vivo* Ca^2+^ imaging and data analysis

Glomerular imaging was performed using a VNO-AOB *ex vivo* preparation (Hammen et al., 2014; Meeks and Holy, 2009). Briefly, after deep isofluorane anesthesia, mice were decapitated and placed into ice-cold aCSF. The anterior skull including the snout was dissected from the skull, and then hemisphered along the midline. The single hemisphere of the mouse snout including the olfactory bulb was adhered to a small plastic plank and submerged in rapidly circulating aCSF in a custom dissection chamber. Under a dissection microscope, the septal cartilage and bone overlying the vomeronasal nerve was carefully removed, the AOB separated from the frontal neocortex, and a 0.0045” OD polyimide cannula (A-M Systems, Carlsborg, WA, USA) was then placed into the VNO for ligand delivery. Volumetric images of the AOB were then acquired using OCPI microscopy using the same stimulus protocols described for VNO imaging. Maps of volumetric glomerular activation were generated as described in (Hammen et al., 2014)(**Supplementary Fig. 5**).

### VNO slicing and single VSN collection

For FLICCR-seq experiments, acute VNO slices from OMPxAi96 mice were prepared on a vibrating microtome (Model VT1200S, Leica) and placed into a modified slice chamber (Warner Instruments). An upright electrophysiological rig based on the Nikon FN1 housing was modified to accommodate a custom computer-controlled stimuli delivery system. At least three randomized, interleaved trials of BA stimuli (Ringer’s, 1μM CA, 1 μM DCA, 1 μM CDCA, 1 μM LCA) were applied to the tissues. A heat map representation of the change in fluorescence intensity (ΔF/F) was generated in near-real-time by a customized MATLAB script to identify reliably responsive VSNs in the tissue. A modified patch pipette (tip diameter 1.5 - 2 µm) was placed near responsive neurons by comparing the ΔF/F image to infrared differential interference contrast (IR-DIC) images. Verification of the pipette placement near the intended VSN was performed by pulsing the tissue with BA stimuli while the pipette was within 5 µm of the cell of interest. Once the location of the BA-responsive VSN was confirmed, the VSN soma was gently aspirated into the pipette tip, and then placed into RNA preservation buffer for later single-cell RNA sequencing. Throughout the single cell selection procedure, images and movies were taken of the cellular and pipette positions, and careful notes taken about the responsiveness of the VSN, whether any unintended material (*e.g*., an adjacent dead cell on the outside of the pipette) was visible. mRNA from each individual VSN sample was converted to cDNA using the SMART-Seq v4 Ultra Low Input RNA Kit (Takara Bio USA, Inc.). The quality of each individual cDNA sample was assessed using an Agilent TapeStation, and samples lacking cDNA fragments spanning the 500-2500 bp size were excluded from further processing and analysis. cDNA libraries were then prepared using Nextera XT DNA Library Prep Kit (Illumina, CA, USA), evaluated again for fragment size using the Agilent TapeStation. Each individual sample cDNA library was barcoded using Nextera XT Index kit (Illumina), and diluted in order to approximately match concentrations across samples. Indexed sample libraries were pooled and sequenced on the MiSeq platform (150 bp, paired end, ∼12M reads/pooled library) by the staff of the McDermott Next Generation Sequencing Core at UT Southwestern Medical Center.

### Single cell RNAseq analysis

Individual fastq files for each VSN sample were initially generated from raw data by bioinformaticians in the McDermott Next Generation Sequencing Core. Samples were trimmed via Trimmomatic (Bolger et al., 2014) and aligned to the mm10 *mus musculus* reference genome using the STAR aligner (Dobin et al., 2013). A total of 158 samples were analyzed, including 145 single VSN samples, 7 bulk VNO tissue samples, and 6 kit controls. The average number of mapped reads per sample was 90,636 (median 83,955). A matrix of raw counts per gene (using HTseq, ENSEMBL version 90) for each sample was generated and imported into Seurat v 2.3.4 (Butler et al., 2018) for subsequent analysis. Briefly, the raw reads were normalized using the “lognormalize” subroutine, the top differentially expressed genes were selected using the “FindVariableGenes” function, and the resulting normalized data scaled using the “ScaleData” function. Principal components analysis (PCA) was run on the variable genes, and approximately 10 PCs were selected that contained the most useful gene expression eigenvectors. Clustering was performed using the default routines of the “FindClusters” function (“resolution” value 1.0), and data were prepared for visualization using UMAP (McInnes et al., 2018). Differential gene expression analysis comparing BA-responsive and BA-unresponsive populations was conducted using the Wilcoxon-Mann-Whitney test (via the “FindMarkers” function in Seurat). For this comparison the log fold-change threshold was set at 0.5 and the minimum percentage of expressing samples set to 0.01 (*i.e*. at least 1 sample). The differentially expressed genes with unadjusted differential expression p-values less than 0.001 were sorted by log fold-change (Tables 1 and 2).

**Table 1.**
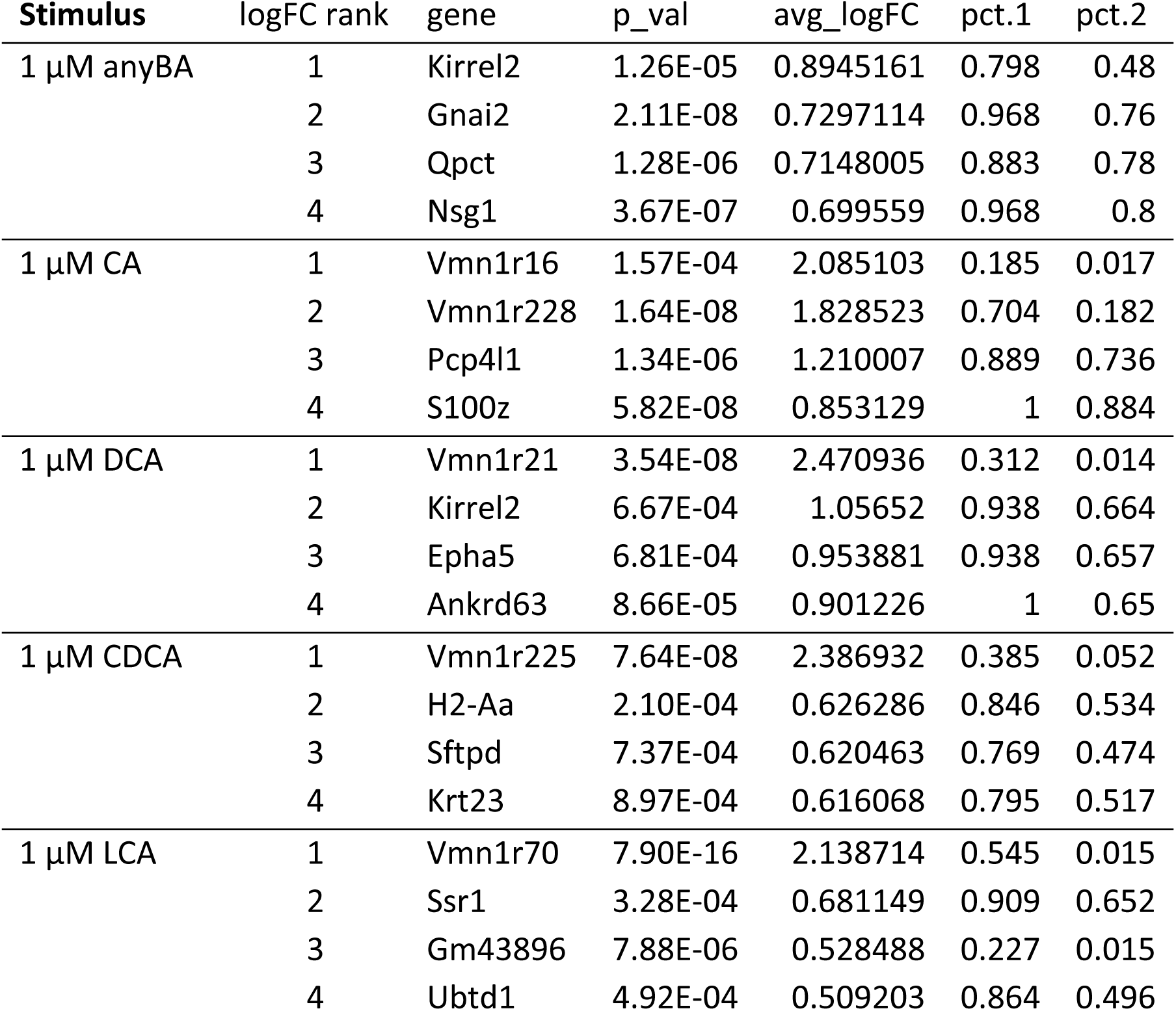
Identification of enriched genes in VSNs, grouped by bile acid sensitivity. Differential expression across all genes was evaluated by the Wilcoxon test, with all genes with p-values less than 0.001 considered. Accepted genes were then rank-ordered based on their average log fold-change (logFC) values.

**Table 2.** Binomial probability estimates of bile acid sensitivity, grouped by gene expression. VSNs found to express a given gene were grouped, and their bile acid sensitivities tested against expectations given the binomial distribution. For example, 5 of the 7 cells expressing Vmn1r16 (71%) responded to 1 µM cholic acid (CA). Of the 158 total samples in the dataset, the number of 1 µM CA-responsive VSNs was 27 (17%). Based on the binomial distribution, the probability of observing 5 or more CA responsive VSNs in the Vmn1r16-expressing set is 0.00219.

| Stim. | *Vmn1r16* | *Vmn1r21* | *Vmn1r70* | *Vmn1r89* | *Vmn1r225* | *Vmn1r228* | *Vmn1r5* | *Vmn1r82* | *Vmn2r1* | *Vmn2r2* | *Vmn2r54* | *Vmn2r89* | *Vmn2r110* | *Gnai2* | *Gnao1* | *Egr1* | *Fos* | *Xntrpc* | *Gapdh* |
| --- | --- | --- | --- | --- | --- | --- | --- | --- | --- | --- | --- | --- | --- | --- | --- | --- | --- | --- | --- |
| anyBA | 0.401 | 0.220 | 0.035 | 0.661 | 0.004 | 0.004 | 1.000 | 1.000 | 1.000 | 1.000 | 0.895 | 0.960 | 0.064 | 0.017 | 1.000 | 0.279 | 0.137 | 0.100 | 0.383 |
| CA | 0.002 | 1.000 | 1.000 | 0.717 | 1.000 | 4.01E-05 | 1.000 | 1.000 | 0.822 | 0.980 | 0.339 | 1.000 | 1.06 E-4 | 0.246 | 0.944 | 0.175 | 0.381 | 0.169 | 0.488 |
| DCA | 0.151 | 5.67E-05 | 1.000 | 0.892 | 1.000 | 0.073 | 1.000 | 1.000 | 0.732 | 0.892 | 1.000 | 1.000 | 0.054 | 0.263 | 0.870 | 0.028 | 0.299 | 0.257 | 0.550 |
| CDCA | 1.000 | 1.000 | 0.981 | 0.614 | 9.56E-07 | 0.176 | 1.000 | 1.000 | 0.999 | 0.997 | 1.000 | 0.298 | 0.921 | 0.119 | 1.000 | 0.694 | 0.185 | 0.515 | 0.440 |
| LCA | 1.000 | 1.000 | 8.63E-08 | 0.571 | 0.500 | 0.996 | 1.000 | 1.000 | 0.692 | 0.956 | 0.647 | 0.893 | 1 | 0.381 | 0.959 | 0.561 | 0.514 | 0.281 | 0.513 |

For additional analysis of gene expression patterns within the vomeronasal gene family, a subset of the log-normalized data including gene expression information for VRs, FPRs, and selected housekeeping and VNO-associated genes (including *Actb, Gapdh, Gnai2, Gnao1, Gnaq, Gnas, Obp2a, Obp2b, Omp, Osbp, Osbp2, Slc9a3r1, Trpc2*) were subjected to an independent cluster analysis using previously-reported routines based on the “mean-shift” method (Comaniciu and Meer, 2002; Meeks et al., 2010). Clusters identified using this alternative method were organized for display purposes and the binomial distribution used to quantify the likelihood of specific clusters containing the observed number of bile acid-responsive VSNs.

### *In vivo* BA exposure and *in situ* hybridization

*In situ* hybridization was performed on VNO slices obtained from 24 C57Bl/6J mice (12 males and 12 females). Briefly, mice were lightly anesthetized via isofluorane and 20 µL of one of the follow stimuli was pipetted directly onto the mouse snout (∼10 µL per nostril); Ringer’s, 1 mM BA mix, 100 μM BA mix, or female mouse urine (not shown). Mice were returned to clean cages and perfused with 4% paraformaldehyde 30 minutes following exposure. The VNOs were extracted and later sectioned using a cryostat (Model VT1200s, Leica, Buffalo Grove, IL) and mounted onto slides. Custom BaseScope probes for *Vmn1r16, Vmn1r21, Vmn1r70, Vmn1r225, Vmn1r228*, and *Vmn2r54* were generated by a commercial vendor (Advanced Cell Diagnostics; Newark, CA). A BaseScope probe for the immediate-early gene Egr1 was purchased, and both probes independently marked via BaseScope Duplex Detection Reagent Kit (Advanced Cell Diagnostics; Newark, CA) following the manufacturer’s instructions. The samples were counterstained stained with dilute hematoxylin/eosin and slides imaged using a Nanozoomer S360 slide scanner (Hamamatsu, Japan) maintained by the UT Southwestern Whole Brain Microscopy Facility.

### *In situ* hybridization quantification

Nanozoomer images were exported to high-resolution color tiff images and then processed using publicly available image classification software, ilastik. Briefly, the “Pixel Classification” plugin for ilastik, which implements voxel-wise, supervised random forest machine learning, was used to produce probability map images of the positions of VR- and *Egr1*-positive puncta. Representative blocks of tissue sections for each VR probe were used as model training datasets, and the same model was applied to 2,748 VNO tissue sections across 210 tissue blocks derived from 24 experimental animals. The probability map images were then used as the input images for CellProfiler, which was used to segment *Egr1-* and VR-positive areas. Finally, the output images from CellProfiler were assembled via custom a MATLAB user interface that allowed users to draw and annotate regions of interest for each imaged vomeronasal epithelium (*e.g*., sex, treatment condition). Additional MATLAB scripts were used to assemble the data and perform statistical comparisons. The block-wise design of our section mounting strategy allowed unambiguous assignment of tissues to their proper treatment condition and sex, but did not allow us to unambiguously assign each section to a specific animal. We pooled sections by condition and sex, then performed statistical comparisons using the Kruskal-Wallis Test followed by post-hoc multiple comparisons tests as appropriate.

### Data availability

Raw gene expression data have been deposited on [REPOSITORY] and all analysis code is available from the corresponding author upon request.

## Results

### Sensitive and selective VSN responses to bile acids

BAs are naturally-occurring AOS ligands that are found abundantly in mouse and other animal feces (Doyle et al., 2016; Hofmann et al., 2010). However, the sensitivity and selectivity of VSNs to BAs has not been deeply explored. In order to quantify VSN BA tuning we performed population VSN Ca^2+^ imaging via OCPI microscopy (**Fig. 1A**). We began by stimulating VSNs with a concentration series of two closely related monomolecular BAs at concentrations spanning 100 nM to 10 µM (**Fig. 1B**). Monomolecular BA ligands elicited reliable, concentration-dependent responses from VSNs across multiple randomized, interleaved trials in this preparation (**Fig. 1C, D**). We utilized both GCaMP6s- and GCaMP6f-expressing preparations (VNO epithelia from both OMPxAi96 and OMPxAi148 mice), which produced comparable results (data not shown), and therefore present the responses of both throughout this manuscript.

**Figure 1:**
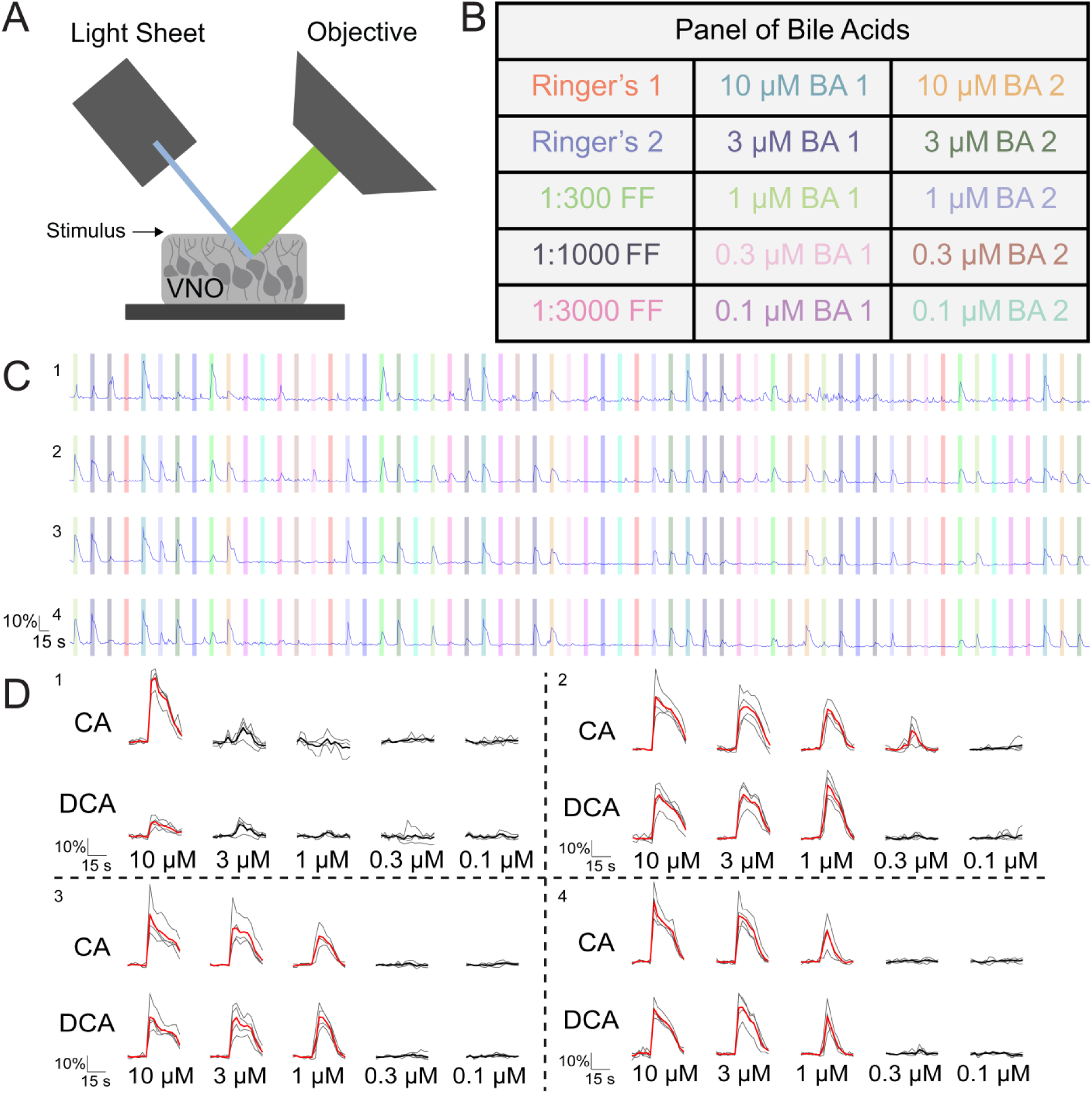
Evaluating vomeronasal sensory neuron responses to monomolecular bile acids with population VSN Ca^2+^ imaging via OCPI microscopy. (**A**) OCPI (light sheet) microscopy imaging set up. Thousands of VSNs in the intact vomeronasal epithelium were imaged via this setup to measure the responses across a panel of ligands. (**B**) Experimental stimulus panel consisting for 5 concentrations of two bile acids (BAs) and positive (dilute female mouse feces) and negative (Ringer’s) controls. (**C**) Representative fluorescence intensity traces of 4 individual VSNs over the course of one experiment. Colored bars represent ligand presentation corresponding to ligands depicted in (**B**). Each stimulus presentation lasted 5 stacks (∼15 s), with 15 stacks (∼45 s) between stimulus presentations. (**D**) Representative across-trial responses of the same VSNs in (**C**). The bolded red trace shows the mean response across all trails. FF, Female Feces; BA, Bile Acids; CA, Cholic Acid; DCA, Deoxycholic Acid

From these datasets, we first performed a pair-wise comparison between VSN responses to a primary bile acid, cholic acid (CA), and its gut bacteria-dependent derivative deoxycholic acid (DCA). These molecules differ from each other only by the presence or absence of a hydroxyl group on the 7-carbon (**Fig. 2**). We observed that many VSNs reliably responded to one or both of these ligands **Fig. 2A, B**). Across the imaged VSNs we observed populations of neurons which responded selectively to CA only (16.01%), DCA only (16.58%), as well as a population that responded to both CA and DCA (20.21%, **Fig. 2C**). As a control for possible nonselective effects of BAs on VSN Ca^2+^ signaling, and relatively high rates of VSN spontaneous activity (**Supplementary Movie S1**), in this and every other experiment we analyzed responses of VSNs that demonstrated a visible change in GCaMP6 fluorescence to at least one stimulus in the panel (by inspecting single-trial ΔF/F). We found that cells chosen based on single-trial (presumably spontaneous) responses were not reliably responsive to either CA or DCA (47.2%), suggesting that BAs do not cause nonselective VSN Ca^2+^ increases. Within the subset of VSNs that responded selectively to CA, a substantial subset (7.7%, representing 1.2% of all evaluated VSNs) were sensitive to CA at 0.1-0.3 µM (**Fig. 2C**), indicating that – for this comparison – these neurons were CA specialists. The majority of neurons that responded reliably to both CA and DCA were sensitive to both ligands at the 1 µM range (74.7%, representing 15.1% of all evaluated VSNs). Others in this category were only responsive at the 3-10 µM range (25.3%, or 5.1% overall). Thus, it was clear that VSNs, both individually and as a population, can distinguish between these functionally and structurally similar BAs.

**Figure 2:**
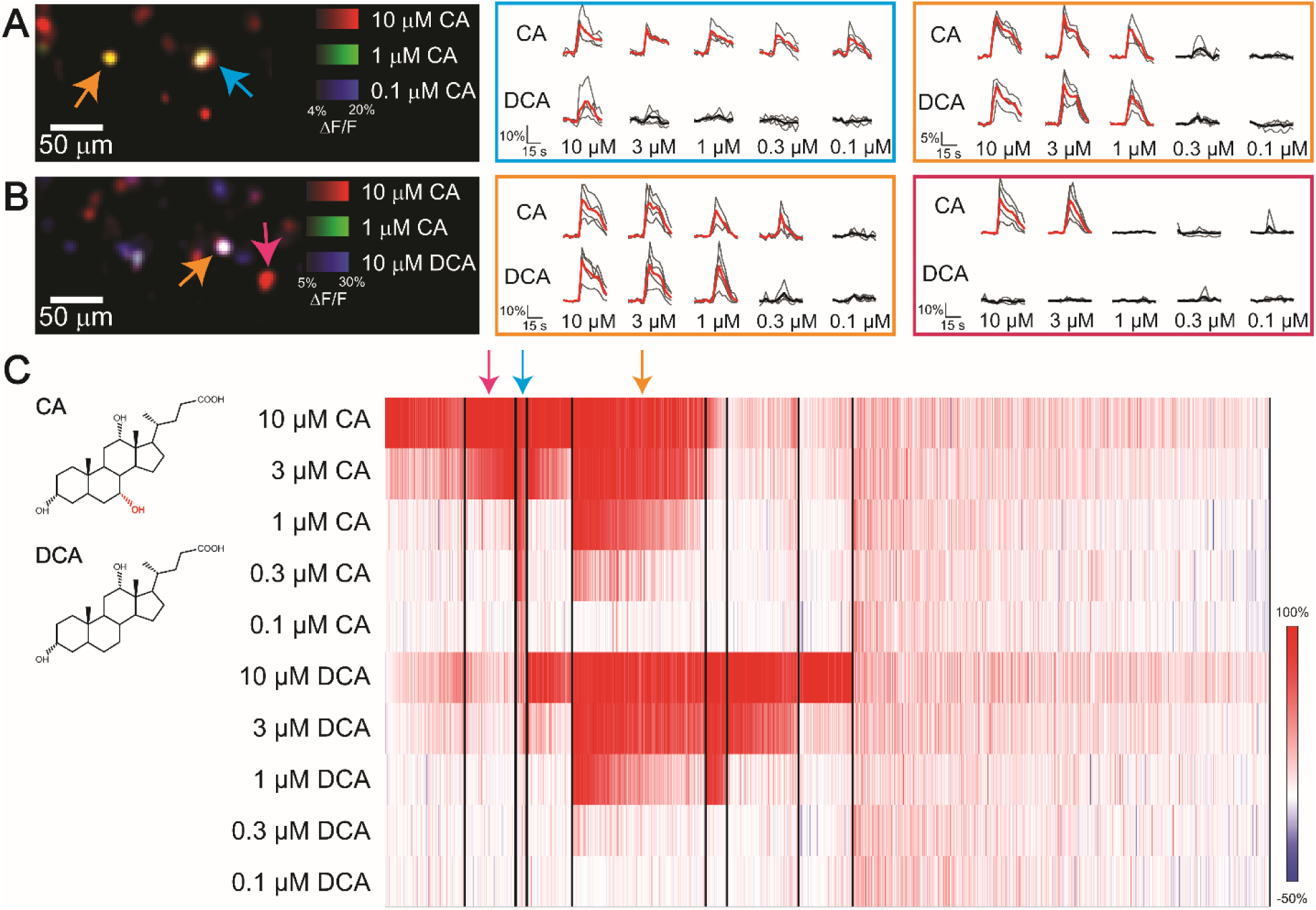
Vomeronasal sensory neuron responses to cholic acid and deoxycholic acid. (**A-B**) Representative colorized images of vomeronasal sensory neurons response (ΔF/F) in a single frame of and OCPI image stack (left). The responses to 10 µM CA (red), 1 µM CA (green), and 0.1 µM CA (blue) are shown in (**A**). The responses of VSNs to 10 µM CA (red), 1 µM CA (green), and 10 µM DCA (blue) are shown in (**B**). Across-trial VSN responses are plotted as individual traces. Bolded trace indicates the mean response across all stimulus repeats. Responses in the cyan box correspond to the VSN indicated by the cyan arrow, those in the orange box correspond to those indicated by orange arrows, and traces in the magenta box correspond to the neuron indicated by the magenta arrow. (**C**) Clustered heat map of VSN response to CA and DCA (structures to left, with red annotations indicating key differences) at varying concentrations. Each column indicates an individual neuronal response. Cluster divisions are indicated by black vertical lines. Arrows above the heat map highlight the clusters in which neurons from (**A**) and (**B**) fall. Shown are the responses of 1,222 VSNs from 3 OMPxAi96 animals. All experiments included at least 3 randomized, interleaved stimulus repeats.

CA and DCA are both found in mouse feces and are closely related in structure, but previous work indicated that mouse VSNs are also sensitive to lithocholic acid (LCA), a BA that is derived from a different primary BA (chenodeoxycholic acid, CDCA), and which is not present at detectable levels in mouse feces (Doyle et al., 2016; Hofmann et al., 2010) (**Fig. 3**). In separate experiments, we examined the response profiles of VSNs to CA and LCA across the same 100 nM – 10 µM concentration range (**Fig. 3**). Confirming earlier studies in the downstream AOB (Doyle et al., 2016), we observed reliable VSN responses to LCA (**Fig. 3A, B**). Clustering based off VSN response profiles revealed populations of neurons that responded selectively to CA only (21.5%), LCA only (14.8%), and both CA and LCA (11.0%, **Fig. 3C**). As before, a subset of CA-selective VSNs showed sensitivity to CA in the 0.1-0.3 µM range (2.1%, 0.45% overall, **Fig. 3C**). We noted that many VSNs that were analyzed based on spurious single-trial (likely spontaneous) activity showed substantial across-trial mean ΔF/F responses to 10 µM LCA (**Fig. 2C**). LCA can be cytotoxic (Katona et al., 2009), so the increased mean ΔF/F to 10 µM LCA in some VSNs may reflect mild cytotoxicity. Each of these pair-wise concentration-response experiments produced similar results: there appeared to be BA specialists and generalists with submicromolar-to-micromolar BA sensitivities. For completeness, we performed pairwise concentration-response experiments for the four major BAs in this study, which included CA, CDCA, deoxycholic acid (DCA), and LCA (**Supplementary Figs. 1-4**). Each comparison revealed nuanced differences (*e.g*., CA and CDCA, two primary bile acids, showed the highest degree of VSN co-activation, **Supplementary Fig. 1**), but in every pairwise comparison we found evidence of BA specificity and selectivity. Cumulatively, these data indicate that, as we previously hypothesized, there are likely to be many BA-sensitive receptors expressed by VSNs.

**Figure 3:**
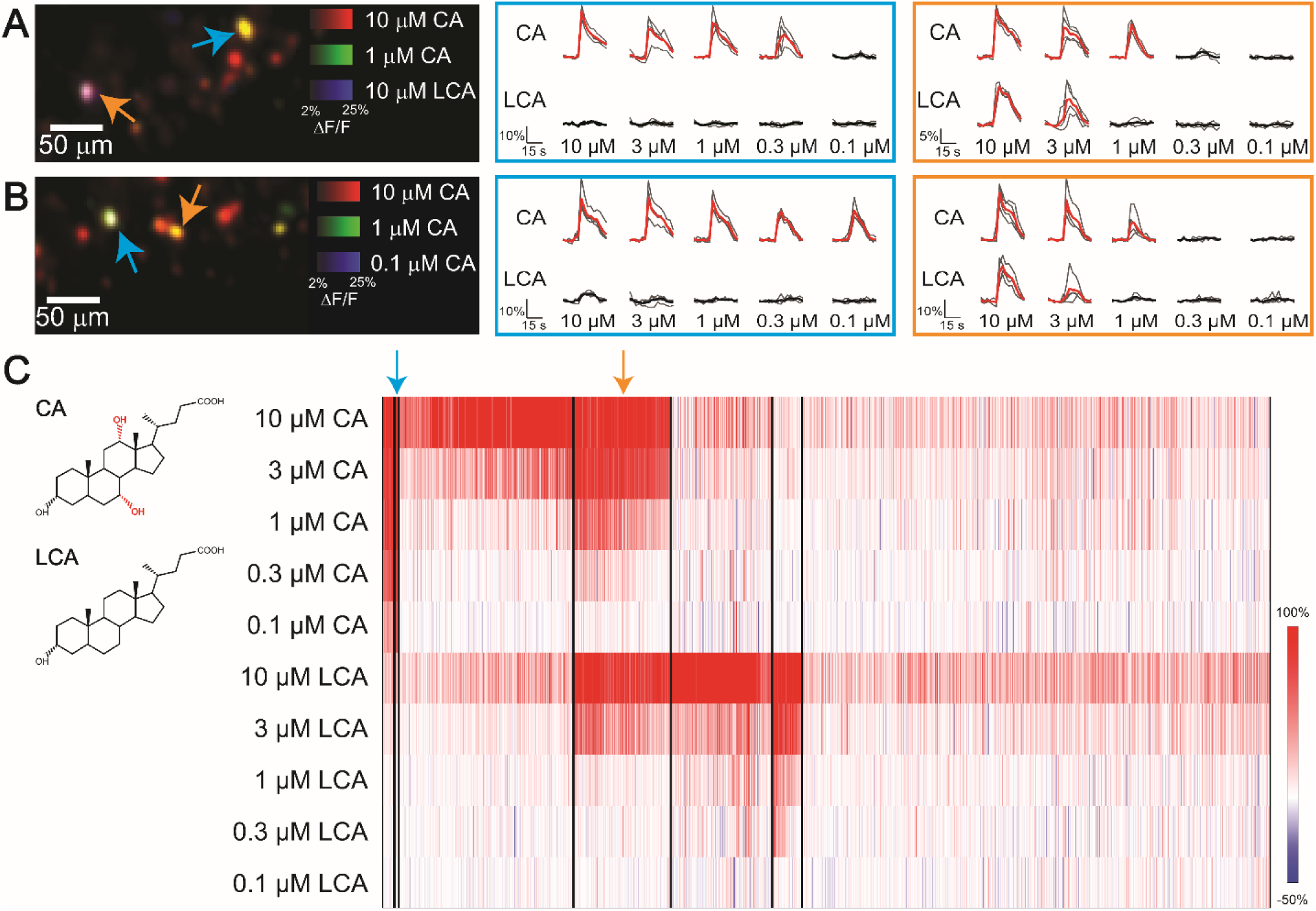
Vomeronasal sensory neuron responses to cholic acid and lithocholic acid. (**A-B**) Representative colorized images of vomeronasal sensory neuron responses (ΔF/F) in a single frame of an OCPI image stack (left). The responses to 10 µM CA (red), 1 µM CA (green), and 10 µM LCA (blue) are shown in (**A**). The responses of VSNs to 10 µM CA (red), 1 µM CA (green), and 0.1 µM CA (blue) are shown in (**B**). Across-trial VSN responses are plotted as individual traces. Bolded trace indicates the mean response across all stimulus repeats. Responses in the cyan box corresponds to the VSN indicated by the cyan arrow while those in the orange box corresponds to those indicated by orange arrows. (**C**) Clustered heat map of VSN response to CA and LCA (structure at left, with red annotations indicating key differences) at varying concentrations. Each column indicates an individual neuronal response. Cluster divisions are indicated by black vertical lines. Arrows above the heat map highlight the clusters in which neurons from (**A**) and (**B**) fall. Shown are the responses of 1,362 VSNs from 3 OMPxAi96 animals. All experiments included at least 3 randomized, interleaved stimulus repeats.

### VSNs shown broad responses to bile acids and sulfated steroids

The high amount of overlap in VSN sensitivities seen in pair-wise BA tuning comparisons suggested that a different, broader assay would be required to determine whether, say, 3 or 30 different VSN populations (and presumably the VRs they express) were sensitive to these 4 common BAs. Moreover, questions remain about whether BA-sensitive VSNs are also activated by other steroid ligands (Fu et al., 2015; Haga-Yamanaka et al., 2014; Nodari et al., 2008; Xu et al., 2016). We conducted further Ca^2+^ imaging experiments in which we exposed the VSNs to a broad panel of BAs, including CA, DCA, CDCA and LCA (each at 1 µM and 10 µM) and four sulfated steroids (each at 10µM). This panel of sulfated steroids included an androgen (A6940), an estrogen (E0893), a pregnanolone (P3865) and a glucocorticoid (Q1570), each of which has been shown to activate specific subsets of VSNs (Meeks et al., 2010; Turaga and Holy, 2012)(**Fig. 4**). Again, VSNs showed reliable across-trial responses to this panel of ligands (**Fig. 4A, B**).

**Figure 4:**
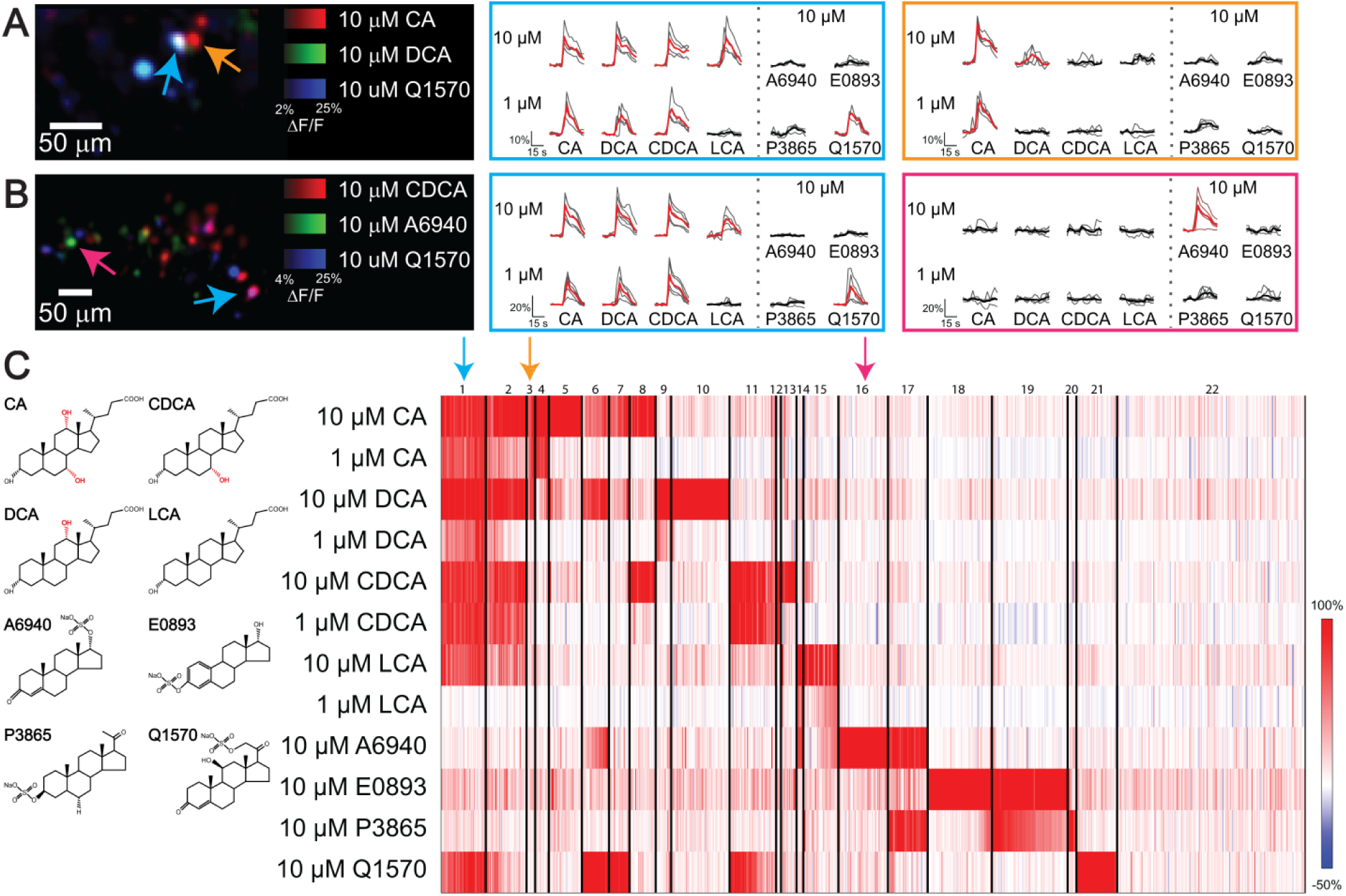
VSN responses to a panel of monomolecular ligands and sulfated steroids. (**A-B**) Representative colorized images of VSN responses (ΔF/F) in a single frame of an OCPI image stack (left). The responses to 10 µM CA (red), 10 µM DCA (green), and 10 µM Q1570 (blue) are shown in (**A**). The responses of VSNs to 10 µM CDCA (red), 10 µM A6940 (green), and 10 µM Q1570 (blue) are shown in (**B**). Across-trial VSN responses are plotted as individual traces. Bolded trace indicates the mean response across all stimulus repeats. Responses in the cyan box correspond to the VSN indicated by the cyan arrow, those in the orange box correspond to those indicated by orange arrows, and traces in the magenta box correspond to the neuron indicated by the magenta arrow. (**C**) Clustered heat map of VSN response to a panel of monomolecular ligands. Each ligand is shown at left to highlight structural differences, with red annotations indicating key differences among bile acids. Each column indicates an individual neuronal response. Cluster divisions are indicated by black vertical lines. Arrows above the heat map highlight the clusters in which neurons from (**A**) and (**B**) fall. Shown are the responses of 890 VSNs from 3 OMPxAi96 animals. All experiments included at least 3 randomized, interleaved stimulus repeats.

Cluster analysis of VSN tuning properties to this broad BA and sulfated steroid panel revealed several interesting features (**Fig. 4C**). First, we found that a substantial population of VSNs (**Fig. 4C**, Clusters 1 & 2) were sensitive to all 4 BAs in this panel at 10 µM, indicating that many of the VSNs that were co-activated in pairwise comparisons are members of an extremely broadly BA tuned VSN population. Within this broadly BA-tuned group of VSNs were cells that were also activated by 10 µM of the sulfated glucocorticoid Q1570 (**Fig. 4C**, Cluster 1). This experiment revealed additional evidence of BA sensitive and selective VSN subpopulations (**Fig. 4C**, Clusters 3-15), and identified many previously-studied sulfated steroid-tuned VSN subpopulations (**Fig. 4C**, Clusters 16-21)(Lee et al., 2019; Meeks et al., 2010; Turaga and Holy, 2012; Xu et al., 2016). Also noteworthy were Clusters 6, 7, 11, and 14, which showed some degree of co-activation by BAs and sulfated steroid ligands. Overall, we found that 29.1% of analyzed VSNs were activated by at least one BA in this panel, 32.3% were activated by at least one sulfated steroid, and 16.9% were activated by at least one BA and one sulfated steroid. The specific patterns of BA and sulfated steroid activity indicate, at a minimum, that there are dozens of VSN populations that encode BA identity. Moreover, the emergence of several BA- and sulfated steroid-sensitive populations indicates that VSNs utilize receptors (presumably VRs) that are sensitive to steroids with substantially different chemical features and are derived from distinct biochemical pathways in the emitter.

Previous studies have determined that steroid-sensitive VSNs express members of the V1R subfamily, which happen to project axons selectively to the anterior subregion of the AOB (Haga-Yamanaka et al., 2014; Isogai et al., 2011; Lee et al., 2019; Meeks et al., 2010). Based on the superficial chemical similarities between BAs and other steroids, and the partial overlap seen between BA- and sulfated steroid responsive VSNs, we hypothesized that V1Rs are likely serving as BA receptors. As a first pass at identifying the BA-sensitive VR family, we performed GCaMP6f Ca^2+^ imaging in specialized *ex vivo* preparations that maintain functional connections between the VNO and AOB (Hammen et al., 2014). Using the same panel of BA and SS ligands shown in **Figure 4**, we stimulated the VNO in these *ex vivo* preparations while performing volumetric imaging of the AOB glomerular layer (where VSN axons terminate, **Supplementary Fig. 5**). Consistent with our hypothesis of V1R BA sensitivity, we observed glomerular layer activation in small, distributed regions of interest resembling individual glomeruli in the anterior AOB (**Supplementary Fig. 5**). Across these experiments we did not identify a distinct spatial region or evidence of spatial co-localization between specific BA-tuned glomerular subpopulations (Hammen et al., 2014), but more detailed investigations will be needed to fully evaluate the spatial distribution of steroid- and BA-sensitive glomeruli. Still, these data provide additional evidence that BA-sensitive VSNs express members of the V1R family.

### Function-forward BA receptor identification via FLICCR-seq

The collective evidence of sensitive, selective, and diverse VSN BA tuning indicated that many BA-sensitive receptors are expressed by BA-sensitive VSNs. Identifying ligand-receptor interactions in the VNO remains a challenging task. We developed FLICCR-seq, a function-forward approach that uses GCaMP6 physiology at the front end, to identify and isolate BA-sensitive VSNs prior to single cell RNAseq (scRNAseq, Methods, **Figure 5A**). We placed acute VNO slices from OMPxAi96 mice into the recording chamber of a modified slice electrophysiology rig. Based on OCPI Ca^2+^ imaging results, we selected 1 µM as the single BA concentration for stimulating BA-sensitive VSNs in these experiments (**Figs. 2-4**). As in OCPI experiments, we used computer-controlled stimulus delivery to apply no fewer than three randomized, interleaved ligand stimulations to each tissue to identify reliably BA-responsive VSNs (**Fig. 5B, C, Supplementary Movie S2**). After we identified 1 µM BA-sensitive VSNs, we gently aspirated the stimulated cell into a modified patch pipette. The cell and its contents were then processed for scRNAseq. We selected a total of 158 samples, including 145 single VSN samples, 7 bulk VNO samples, and 6 kit controls. Among the 145 single VSN samples were 27 CA-responsive, 16 DCA-responsive, 39 CDCA-responsive, and 22 LCA-responsive cells. We analyzed scRNAseq data using Seurat (Butler et al., 2018), finding gene expression patterns that included a larger, more dispersed group (Clusters 0-2), a smaller, more compact cluster (Cluster 3), and a tiny cluster representing kit (non-VNO) RNA controls (Cluster 4, **Fig. 5D**). Inspecting the expression patterns of VSN-associated transcripts in these clusters revealed that the vast majority of samples in Clusters 0-2 expressed *Gnai2* (encoding the V1R-associated G_αi2_ protein), whereas the vast majority of samples in Cluster 3 expressed *Gnao1* (encoding the V2R-associated G_αo_ protein, **Fig. 5E**). Other genes thought to be expressed by both V1R- and V2R-expressing VSNs, *Trpc2* (encoding the TRPC2 cation channel) and *Ano1* (encoding the anoctamin 1/TMEM16α calcium-activated chloride channel), were found to be expressed by the majority cells in all clusters except kit controls (Cluster 4; **Fig. 5E**).

**Figure 5.**
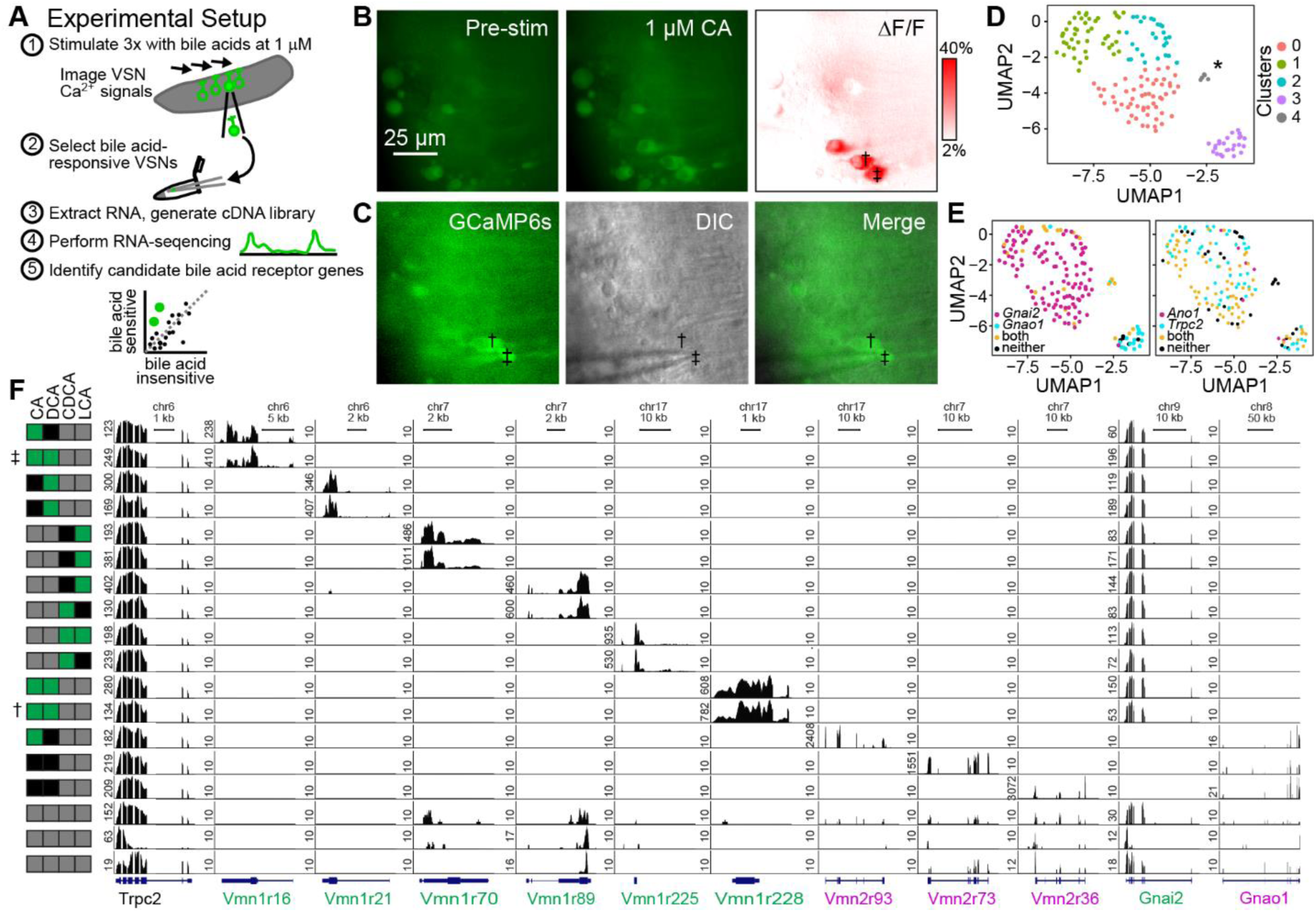
Function-forward selection of BA-sensitive VSNs for scRNAseq. (**A**) Overview of FLICCR-seq experimental setup. VNO slices from OMP-GCaMP6s mice were stimulated with 1 µM BAs a minimum of 3 times. BA-responsive cells are plucked with a modified glass patch pipette and processed for scRNAseq. (**B**) Representative fluorescence images of 1 µM CA-responsive VSNs. The VSNs marked with ‡ and † were selected and processed for scRNAseq. (**C**) Representative images of VSN selection during active stimulation (in this trial with 1 µM DCA). (**D**) UMAP multidimensional scaling of 158 samples, including 145 VSNs collected using these methods. Clusters 0-2 represent subsets of a larger group, whereas Cluster 3 identifies a smaller, isolated cluster. Cluster 4 (grey, asterisk) contained RNA kit control samples, and are accordingly distinct from VSNs. (**E**) *left:* UMAP multidimensional scaling with each cell colorized based on expression of *Gnai2* (G_αi_, magenta) and *Gnao1* (G_αo_, cyan), genes that are associated with V1R-expressing and V2R-expressing VRs, respectively. This analysis indicates that Clusters 0-2 contain V1R-expressing VRs, while Cluster 3 contains mostly V2R-expressing VRs. *right:* Same UMAP scaling with each cell colorized based on the expression of *Ano1* (Anoctamin 1, a.k.a. TMEM16α) and *Trpc2* (cyan), ion channels that have been studied in VSNs. Expression of *Trpc2* or both *Trpc2* and *Ano1* was detected in almost all cells. (**F**) Genome viewer visualization of 18 representative VSNs showing the mapped scRNAseq reads for *Trpc2*, selected V1Rs and V2Rs, and G protein alpha subunits. The rows marked with ‡ and † reflect the cells indicated in (**B**) and (**C**). Note that the number of reads for individual VRs in each VSN was often stronger than for reference genes (*e.g*., *Trpc2, Gapdh*, and *Actb*).

Further inspection of the scRNAseq data for these samples revealed patterns of gene expression consistent with single VSNs, including the near exclusive expression of single VRs (**Fig. 5F, Supplementary Fig. 6**). In addition to the samples containing individual VSNs, we included a small number of VNO samples (including several hundred cells each). In these pooled samples, we detected the expression of many VRs at low expression levels (**Fig. 5F, Supplementary Fig. 6**), indicating that single-VR expression patterns were not an unintended feature of the scRNAseq pipeline, which necessarily involves several amplification steps. Collectively, the depth and quality of scRNAseq data were sufficient for further analysis to attempt to correlate gene expression patterns with patterns of BA responsivity for these VSN samples (**Figs. 5 and 6**).

**Figure 6.**
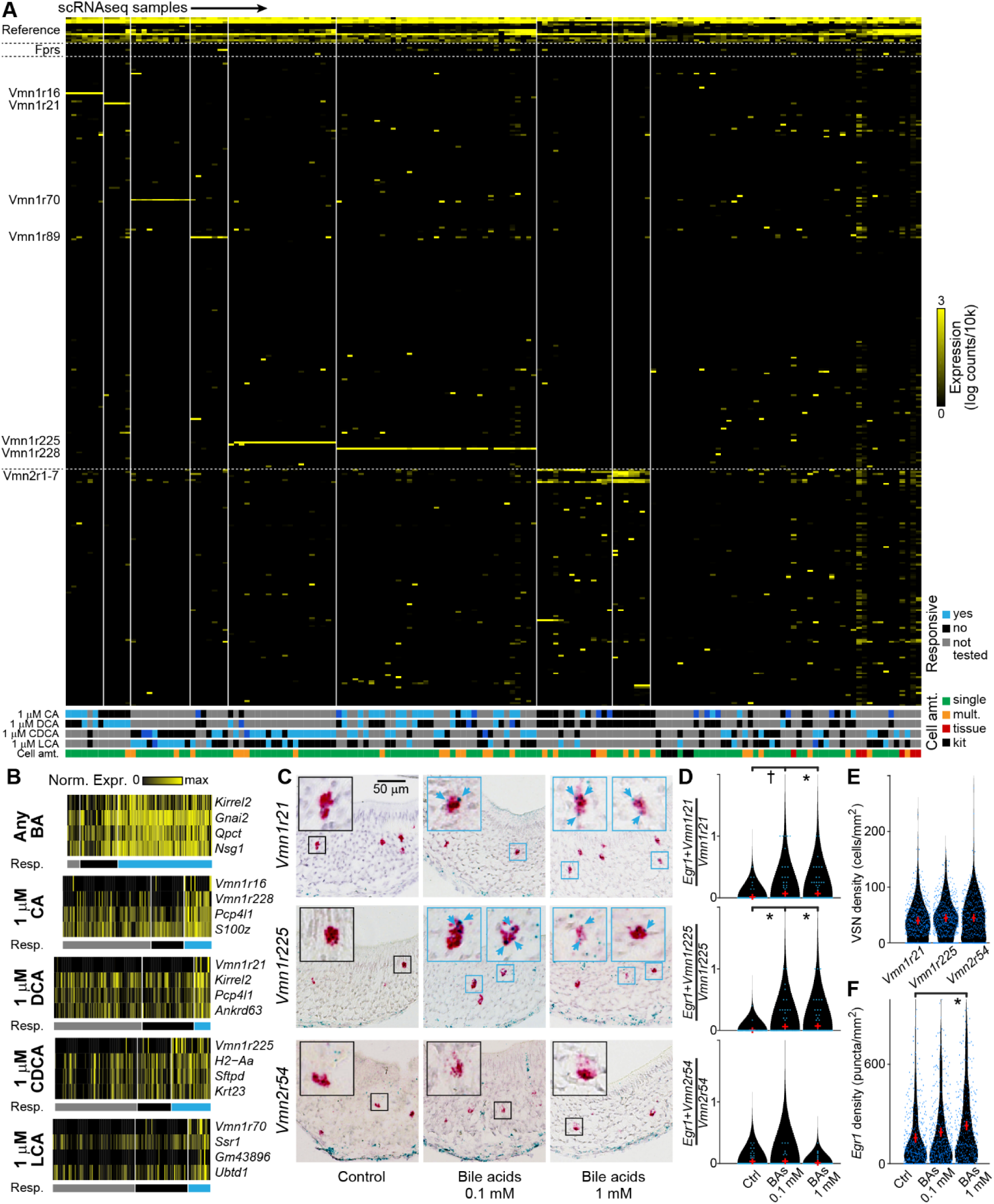
Identification of BA-sensitive vomeronasal receptors. Clustered heat map showing the gene expression for all 158 analyzed samples. Rows are individual transcripts, with divisions between functional subgroups indicated by a horizontal dotted line. At the top are control genes (see Methods). At bottom is a lookup table of responsivity. Cyan indicates that the sample contained a VSN that was reliably responsive to the stimulus, dark blue an inconsistent response, black no response, and gray the ligand was not tested on that VSN. For cell amount (“Cell Amt.”), green indicates a single cell was cleanly picked, orange indicates additional cell material was visible in/on the pipette tip, red indicates a bulk tissue extraction, and black indicates RNA kit control samples. (**B**) Heat map plot of the top 4 differentially-expressed genes for BA-responsive VSNs. In each subplot, the colored horizontal bars indicate the responsivity, with colors representing the same qualities as in **A**. VSNs with unclear responsivity for the specified ligand were excluded from these plots. Expression levels (log counts/10k) were normalized by the maximum values per gene across all samples. (**C**) Duplex RNA *in situ* hybridization micrographs for the immediate-early gene *Egr1* (blue stain) and test vomeronasal receptors (red stain). Colocalized *Egr1* puncta are noted with cyan arrows. Scale bar: 50 µm. Mice were exposed *in vivo* to control saline or a mixture of BAs at 0.1 mM or 1 mM to stimulate VSNs. (**D**) Violin plots of the amount of overlap between *Egr1* staining and each VR observed across multiple VNO slices from 18 mice (3 males and 3 females per condition). † indicates p < 0.1 and **\*** indicates p < 0.05 (Kruskal-Wallis test). (**E**) Violin plot of the density of VR staining across VNO sections from 24 mice (the same 18 from E and 6 additional animals). (**F**) Violin plot of the density of *Egr1* staining across 18 mice (3 males and 3 females per condition). * indicates p < 0.05, †indicates p < 0.1 (Kruskal-Wallis test).

### Identification of five BA-sensitive V1Rs

Inspecting the expression patterns of individual scRNAseq samples revealed that several V1Rs were often expressed by BA-sensitive VSNs (**Fig. 5F, Fig. 6A**). For example, 7 samples expressed *Vmn1r16* (encoding the V1rc29 receptor), 14 samples expressed *Vmn1r70* (encoding the V1rl1 receptor), and 36 samples expressed *Vmn1r228* (encoding the V1re3 receptor, **Fig. 5F**). The apparent enrichment of these VR genes in BA-sensitive channels seemed consistent with a correlation between the expression of these particular V1Rs and BA sensitivity. To determine the probability of achieving these results by chance, we performed differential gene expression analysis on the whole dataset. First, we used responsivity to BAs as a grouping variable (**Fig. 6B, Table 1**). Collapsing all BA-responses regardless of ligand identity (*i.e*., comparing all BA-responsive to all to BA-nonresponsive samples) revealed enrichment of *Kirrel2* and *Gnai2*, which are known to be expressed by V1R-expressing VSNs (Brignall et al., 2018; Prince et al., 2013), but no strong correlation with any individual VR (**Fig. 6B, Table 1**). When we included ligand identity as a grouping variable (*e.g*., comparing all 1 µM CA-responsive to all 1 µM CA-unresponsive samples), this analysis revealed enrichment of 5 individual V1Rs. *Vmn1r16* (V1rc29) and *Vmn1r228* (V1re3) were enriched in 1 µM CA-responsive samples, *Vmn1r21* (V1rc28) was enriched in 1 µM DCA-responsive samples, *Vmn1r225* (V1re5) was enriched in 1 µM CDCA-responsive samples, and *Vmn1r70* (V1rl1) was enriched in 1 µM LCA-responsive samples (**Fig. 6B, Table 1**). We then used a separate approach for evaluating the statistical significance of these enriched receptors, this time by grouping samples based on their expression of the candidate V1Rs (**Fig. 6A, Table 2**). Using this approach, we found that *Vmn1r225-* and *Vmn1r228*-expressing cells were statistically more likely to respond to any BA (regardless of BA identity) than expected by chance, indicating a potential role for these two V1re clade members in broader BA sensitivity (p < 0.01, binomial test, **Fig. 6A, Table 2**). *Vmn1r16*-expressing cells were only enriched in 1 µM CA sensitivity, indicating that *Vmn1r16* (V1rc29) may be a CA-selective receptor (**Fig. 6A, Table 2**). *Vmn1r21*-expressing cells were enriched in 1 µM DCA sensitivity, indicating that *Vmn1r21* (V1rc28) may be a DCA-selective receptor (**Fig. 6A, Table 2**). *Vmn1r70*-expressing cells were enriched in 1 µM LCA sensitivity, indicating that *Vmn1r70* (V1rl1) may be an LCA-selective receptor. Importantly, though many of the samples we collected expressed *Vmn1r89* (V1rj2), which was previously shown to be highly sensitive to certain sulfated estrogens (Haga-Yamanaka et al., 2014), we did not find evidence that *Vmn1r89* or several other expressed receptors (*e.g*., *Vmn2r1, Vmn2r2, Vmn2r54*) were enriched for BA responsiveness (**Table 2**). Collectively, these analyses provide strong evidence for V1R enrichment in populations of VSNs with specific patterns of BA sensitivity, and support the hypothesis that the diverse patterns of BA tuning observed via OCPI microscopy are due to the presence of several BA-sensitive vomeronasal receptors.

The statistical evidence for a correlation between expression of these V1Rs and BA sensitivity patterns was strong, so we sought additional verification of these ligand-receptor pairs. We performed duplex *in situ* hybridization for select VRs (*Vmn1r21, Vmn1r225*, and *Vmn2r54* as a control) and the immediate-early gene *Egr1*, similar to previous studies (Isogai et al., 2011)(**Fig. 6C-E**). We exposed adult male and female mice to a mix of all four of the tested BAs at concentrations of 0.1-1 mM, then performed RNA in situ hybridization (**Fig. 6C**). We observed increased colocalization between *Vmn1r21* and *Egr1* at the 1 mM concentration and increased colocalization between *Vmn1r225* and *Egr1* at both 0.1 mM and 1 mM (**Fig. 6C-D**). In contrast, we did not observe increased colocalization between our negative *Vmn2r54* control and *Egr1* probes (**Fig. 6C-D**). This increased colocalization was not the result of variable densities of *Vmn1r21*+, *Vmn1r225*+, and *Vmn2r54*+ cells (**Fig. 6E**). Despite the relative sparsity of *Egr1+* puncta in these tissues, we did observe increased density of these *Egr1+* puncta in the 1 mM BA-treated animals, and a trend-level increase in *Egr1*+ density in 0.1 mM BA-treated animals (Fig. 6F). These experiments provided additional evidence that these candidate V1Rs are sensitive to BA stimulation *in vivo*.

## Discussion

### VSNs are sensitive, selective BA detectors

Previous studies showed that BAs can elicit robust AOS activation, and the diversity of BA tuning by individual neurons in the AOB (immediately downstream of VSNs) indicated that BAs may be sensed by multiple VSN receptors (Doyle et al., 2016). BAs, which are found abundantly in animal feces, possess several features that could provide useful information to the receiving animal. For example, BAs vary by species, sex, diet, and gut microbiome (Hofmann et al., 2010). Across the animal kingdom, BAs have long been studied as olfactory ligands, and are utilized by several fish species as pheromones (Buchinger et al., 2014). Recent work in zebrafish has identified BA-sensitive ORA-class receptors in the olfactory epithelium (Cong et al., 2019). In the context of our growing knowledge of AOS ligands, BAs are a compelling, but poorly-understood source of mouse social chemosensory information. As such, identifying the mechanisms of detection, projection, and processing of BA information will be important for producing a comprehensive view of the AOS and its many impacts on mammalian social and reproductive behaviors.

Even though BAs were previously shown to reliably elicit AOS activation in the AOB (Doyle et al., 2016) and VNO (Wong et al., 2018), there has been relatively limited information about the sensitivity and selectivity of VSNs to BAs. Other monomolecular AOS steroid ligands have been found to be highly potent VSN activators – some at subnanomolar concentrations – so we sought to thoroughly evaluate VSN BA tuning using OCPI microscopy (**Figs. 1-3, Supplementary Figs. 1-4**). Volumetric Ca^2+^ imaging via OCPI microscopy remains one of the most robust methods for evaluating VSN ligand sensitivity because it enables the unbiased sampling of hundreds to thousands of VSNs per animal (Holekamp et al., 2008; Turaga and Holy, 2012; Wong et al., 2018; Xu et al., 2016). Concentration-response experiments using OCPI in pairs of BAs revealed the presence of BA-specialist and BA-generalist VSNs (**Figs. 1-3, Supplementary Figs. 1-4**). Many of these VSNs showed sub-micromolar BA sensitivity, comparable to other steroid-sensitive VRs (Haga-Yamanaka et al., 2015; Lee et al., 2019).

These experiments show that each of the BAs tested is capable of selectively driving a subset of VSNs. However, some questions remained about whether VNO BA sensitivity was a byproduct of sensitivity to other steroids (*e.g*., polar sex steroids and/or glucocorticoids). Using OCPI microscopy and a broad panel including both BAs and other polar steroids, we found a high degree of segregation between BA-sensitive and BA-insensitive VSNs (**Fig. 4**). This indicates that BA responsiveness is not a general feature of all steroid-sensitive VSNs. It could be the case, however, that the BAs utilized in this panel are not the most potent members of the BA class. Future studies will be needed to determine the full breadth of VNO BA sensitivity, as there are dozens to hundreds of naturally-occurring BA molecules that could have biological relevance in rodents (Hagey et al., 2010).

In the initial experiments describing AOS BA sensitivity, a small number of AOB neurons (downstream of VSNs) demonstrated co-activation by BAs and sulfated glucocorticoids at 10 µM (Doyle et al., 2016). AOB mitral cells innervate multiple AOB glomeruli and are capable of performing excitatory integration (Meeks et al., 2010; Wagner et al., 2006), so it could have been the case that these AOB mitral cells generated these response patterns through excitatory integration of glomerular input from BA- and glucocorticoid-specialist VSNs. Even though we saw largely segregated VSNs sensitivities to BAs and other steroids in this stimulus panel, we identified several VSN populations that were sensitive to both BAs and sulfated glucocorticoids (**Fig. 4**). This shows that some VSNs (and presumably the receptors they express) are capable of cross-class tuning upstream of the AOB circuit. We saw a very small number of VSNs that were co-activated by BAs and sulfated androgen, estrogen, and pregnanolone ligands, but given the relatively narrow focus of this panel, we cannot exclude that other non-BA steroid ligands may co-activate BA-sensitive VSNs. Overall, these studies show that the VSN sensitivity and selectivity for BAs is comparable to most of the other major known AOS ligand classes (Haga-Yamanaka et al., 2015; Kaur et al., 2014; Leinders-Zufall et al., 2000; Nodari et al., 2008; Osakada et al., 2018).

### FLICCR-seq supports the identification of BA-sensitive vomeronasal receptors

An important step toward improving our understanding of mammalian BA chemosensation is to identify the peripheral receptors that endow animals with BA sensitivity. This continues to be a daunting task in the AOS, where limited tools have been available for receptor screening (though see (Dey et al., 2013; Haga-Yamanaka et al., 2014; Lee et al., 2019)). Rather than attempt to recreate heterologous expression systems (Dey et al., 2013; Mainland and Matsunami, 2012; Saito et al., 2004), viral delivery methods (Lee et al., 2019), or transgenic mice for this purpose, we instead sought to develop a method that used primary VSNs in transgenic mice expressing GCaMP6f/s, drawing inspiration from (Haga-Yamanaka et al., 2014). The major drawback in taking this approach is that, assuming proportional expression of the ∼300 known VRs and FPRs expressed by VSNs, less than 1% of the cells in the tissue express any given receptor (creating a “needle in a haystack” problem). We overcame this drawback by developing FLICCR-seq, which enabled us “find the needle in the haystack by making it glow.” In FLICCR-seq, we used clean micropipettes and an epifluorescence slice physiology microscope to pluck BA-responsive VSNs from VNO slices and process them for scRNAseq using established, commercially-available protocols. FLICCR-seq samples were sequenced at relatively high coverage, and with high quality (averaging over 90k mapped reads/sample), resulting in high resolution coverage of expressed genes, including VRs, which are expressed at high relative levels in these cells (**Fig. 5F**). Analysis of FLICCR-seq VSN expression patterns produced strong matches with long-known signatures of V1R- and V2R-expressing cells (*e.g*., *Gnai2, Gnao1*, **Fig. 5**), providing more evidence that FLICCR-seq produces interpretable data. In the context of the growing list of techniques for investigating VSN ligand-receptor interactions, the evidence indicates that FLICCR-seq has several benefits that make it an attractive approach in this difficult-to-study sensory system.

Despite its effectiveness in the context of this study, it is important to note a limitation of our initial implementation of FLICCR-seq, which relies on manual plucking of cells. Unlike droplet-based scRNAseq pipelines (reviewed in Hwang et al., 2018), selecting cells manually from slices introduces the possibility that individual samples may contain the cell of interest plus some unintended material (*e.g*., debris from nearby dead cells). Though practice can reduce these occurrences, completely eliminating such material is nearly impossible. Rather than throw away samples in which a small amount of unintended material was visible, we instead focused on maintaining thorough annotation of each cell picking event, for example by taking continuous screen captures of the procedure and making copious contemporaneous notes (**Supplementary Fig. 6**). Since the goal of FLICCR-seq was to identify the enrichment of receptors in BA-sensitive cells, so long as the identity (and dominant sensory receptor) of any unintentionally collected material was random, we would not expect that contaminating RNA-seq counts would covary with annotated ligand sensitivities. As such, in this context there is little risk of false positives. Even still, improvements in the procedures for cell identification and clean sample picking will make FLICCR-seq even stronger, and will support the discovery of more ligand-receptor matches in the future.

### Identification of five V1Rs sensitive to four common BAs

The FLICCR-seq process allowed us to perform differential expression analysis on BA-sensitive and BA-insensitive VSNs (**Figs. 5-6, Tables 1-2**). Using multiple analytical strategies, five candidate BA-sensitive VRs emerged: *Vmn1r16* (V1rc29), *Vmn1r21* (V1rc28), *Vmn1r70* (V1rl1), *Vmn1r225* (V1re5), and *Vmn1r228* (V1re3). *Vmn1r16* and *Vmn1r21* are both members of the V1rc clade on *mus musculus* chromosome 6 (Rodriguez et al., 2002), and were previously associated with *in vivo* activation by soiled bedding from multiple species (Isogai et al., 2011). These results suggest that components of soiled bedding samples from rodent and non-rodent species activate BA-sensitive VRs. Differential expression analysis revealed enrichment of CA-sensitive cells in *Vmn1r16*-expressing VSN samples (**Fig. 6, Tables 1 and 2**) and DCA-sensitive cells in *Vmn1r21*-expressing samples. These two BA ligands are closely related (DCA is a gut-microbe-dependent metabolite of CA) and are common to many species (Hofmann et al., 2010). Given that we observed CA- and DCA-selective VSNs in OCPI imaging experiments (Figs. 2-4), it seems likely that these two V1rc clade members are expressed by these sensitive, selective VSN populations. For these and other identified BA receptors it is possible that each of the identified receptors is even more sensitive to another natural ligand not present in our panel. However, given that *in vivo* BA exposure causes immediate-early gene induction in *Vmn1r21*-expressing VSNs, it seems likely that these receptors play roles in sensing common BAs in the environment. Similarly, differential FLICCR-seq analysis indicates that *Vmn1r70* (V1rl1)-expressing VSNs are sensitive and selective for LCA a potentially toxic natural metabolite of CDCA that is absent in rodents (**Fig. 6, Tables 1, 2**)(Hofmann et al., 2010). OCPI experiments indicated the presence of VSNs that were sensitive and selective for 1 µM LCA, though this ligand may have caused some nonspecific cytotoxicity at 10 µM (**Figs. 3, 4, Supplementary Figs. 3, 4**). To date there is little known about V1rl1, located on *mus musculus* chromosome 7 adjacent to VRs assigned to the V1re clade (Rodriguez et al., 2002). Our results suggest that V1rl1 may be a selective detector of LCA.

Differential gene expression analysis of *Vmn1r225-* and *Vmn1r228*-expressing samples indicated enrichment for multiple BA-sensitive VSNs (**Fig. 6, Tables 1-2**). *Vmn1r225* (V1re5) and *Vmn1r228* (V1re3) are both located on *mus musculus* chromosome 7 and have been associated with the V1re clade (Rodriguez et al., 2002). These receptors were the most highly enriched in our FLICCR-seq dataset (**Fig. 6**), which was by design enriched for BA-sensitive VSNs. Given this experimental design, one would expect to encounter a higher number of VSNs expressing broadly-tuned receptors than VSNs expressing narrowly-tuned receptors. OCPI analysis, similarly, identified multiple populations of broadly-tuned BA receptors (**Fig. 4**). These results suggest that *Vmn1r225* and *Vmn1r228* encode broad BA receptors. Previous studies indicated that these receptors were activated by animal bedding (Isogai et al., 2011). Interestingly, these studies also indicated that closely-related V1re clade members, *Vmn1r226* (V1re2) and *Vmn1r227* (V1re6), were sensitive to sulfated glucocorticoids (Isogai et al., 2011). Given that OCPI experiments indicated that several classes of broadly BA-tuned VSNs are also sensitive to 10 µM sulfated corticosterone (**Fig. 4**), it seems highly likely that these closely-related receptors are capable of sensing steroids across the BA and glucocorticoid ligand classes.

It is worth noting that we encountered many samples in our FLICCR-seq dataset that expressed *Vmn1r89* (V1rj2), a receptor that has been associated with sulfated estrogen and androgen sensing (**Fig. 6**)(Haga-Yamanaka et al., 2014; Isogai et al., 2011). However, differential gene expression analysis did not indicate enrichment of BA-sensitive VSNs in *Vmn1r89*-expressing FLICCR-seq samples (**Tables 1-2**). It is unclear why our FLICCR-seq dataset included so many *Vmn1r89*-expressing samples. A small number of sulfated estradiol- or testosterone-sensitive VSNs had some BA sensitivity (**Fig. 4**), but these data do not indicate a specific role for this receptor in BA detection.

Taken as a whole, the combination of OCPI and FLICCR-seq experiments produced a deep demonstration of BA sensitivity and selectivity, and identified several V1R-family members that serve as mouse BA receptors. The behavioral impacts of chemosensory BA detection remain unknown, but given the breadth of naturally-occurring BAs and their associations with physiological states like species, sex, diet, and gut microbiome (Hofmann et al., 2010), it seems likely that BAs contribute to large repertoire of AOS-associated behaviors. The idea that BAs could play important roles in vertebrate social/reproductive behaviors is not new, as BAs are known pheromones in lampreys and other fishes (Buchinger et al., 2014). These studies indicate that such roles may also be played by BAs through the AOS in rodents. Overall, these results will support future studies into the molecular mechanisms of BA sensitivity, the network mechanisms that compare and combine BA information with other AOS ligands, and the impacts of BA chemosensation on mouse social behavior.

## Acknowledgements

This work was supported by the National Institute on Deafness and Other Communication Disorders and National Institute of Neurological Disorders and Stroke of the USA National Institutes of Health (Grants R01DC015784 [JPM], R01DC017985 [JPM], R21NS104826 [JPM], and F31DC017661 [WMW]) and the Welch Foundation (Grant number I-1934-20170325 [JPM]). The content is solely the responsibility of the authors and does not necessarily represent the official views of the National Institutes of Health. We thank Natasha Browder and Cara Nielson for technical support, and Dr. Genevieve Konopka and Dr. Fatma Ayhan for critical feedback on the manuscript.

---

**Supplementary Figure 1:**
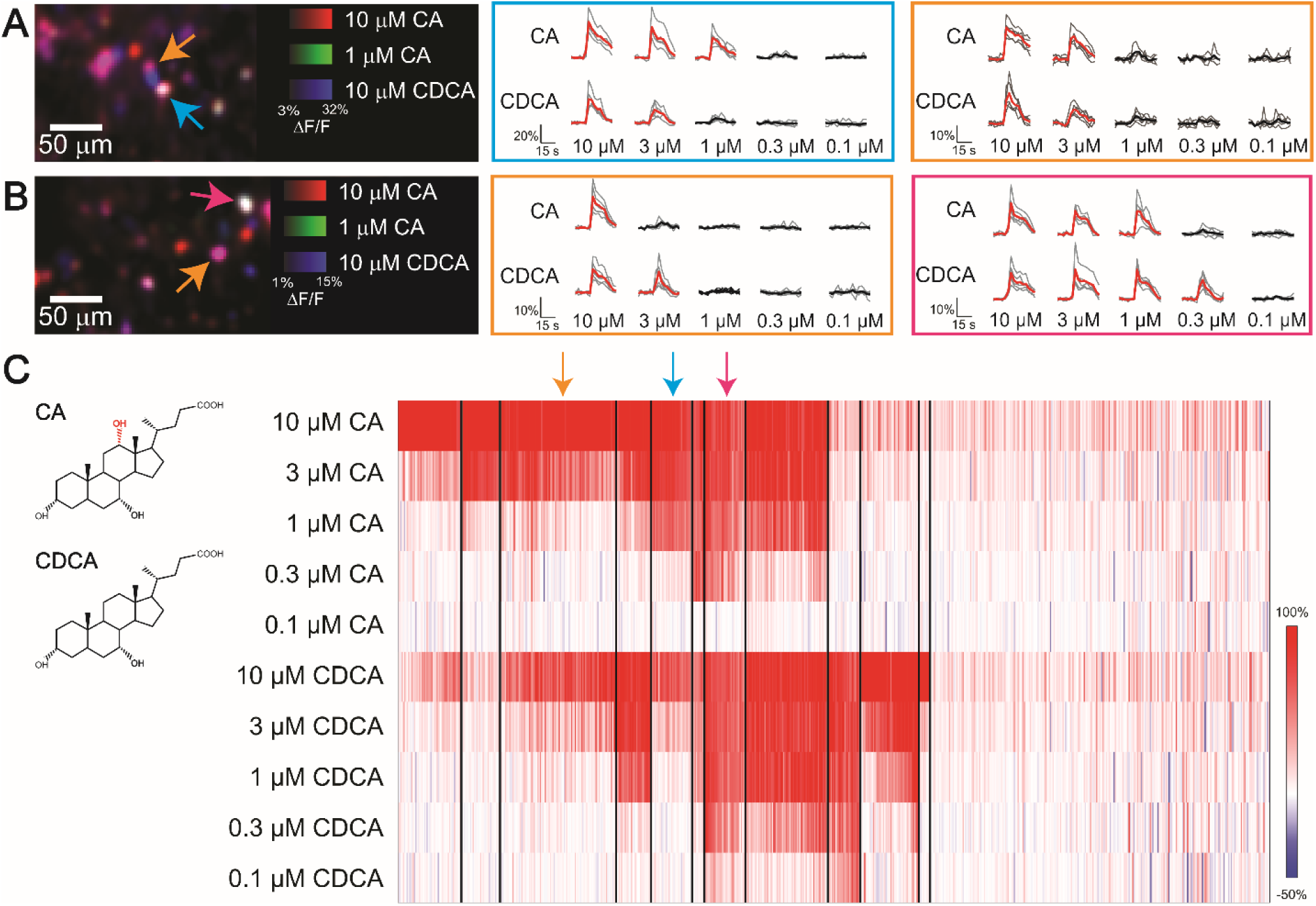
Vomeronasal sensory neuron responses to cholic acid and chenodeoxycholic acid. (**A-B**) Representative colorized images of VSN responses (ΔF/F) in a single frame of an OCPI image stack (left). The responses to 10 µM CA (red), 1 µM CA (green), and 10 µM CDCA (blue) are shown in (**A**). The response of VSNs to 10 µM CA (red), 1 µM CA (green), and 10 µM CDCA (blue) are shown in (**B**). Across-trial VSN responses are plotted as individual traces. Bolded trace indicates the mean response across all stimulus repeats. Responses in the cyan box correspond to the VSN indicated by the cyan arrow, those in the orange box correspond to those indicated by orange arrows, and traces in the magenta box correspond to the neuron indicated by the magenta arrow. (**C**) Clustered heat map of VSN response to CA and CDCA (structures at left) at varying concentrations. Each column indicates an individual neuronal response. Cluster divisions are indicated by black vertical lines. Arrows above the heat map highlight the clusters in which neurons from (**A**) and (**B**) fall. Shown are the responses of 700 VSNs from 3 OMPxAi96 animals (2 males and 1 female). All experiments shown have ≥ 3 repeats each.

**Supplementary Figure 2:**
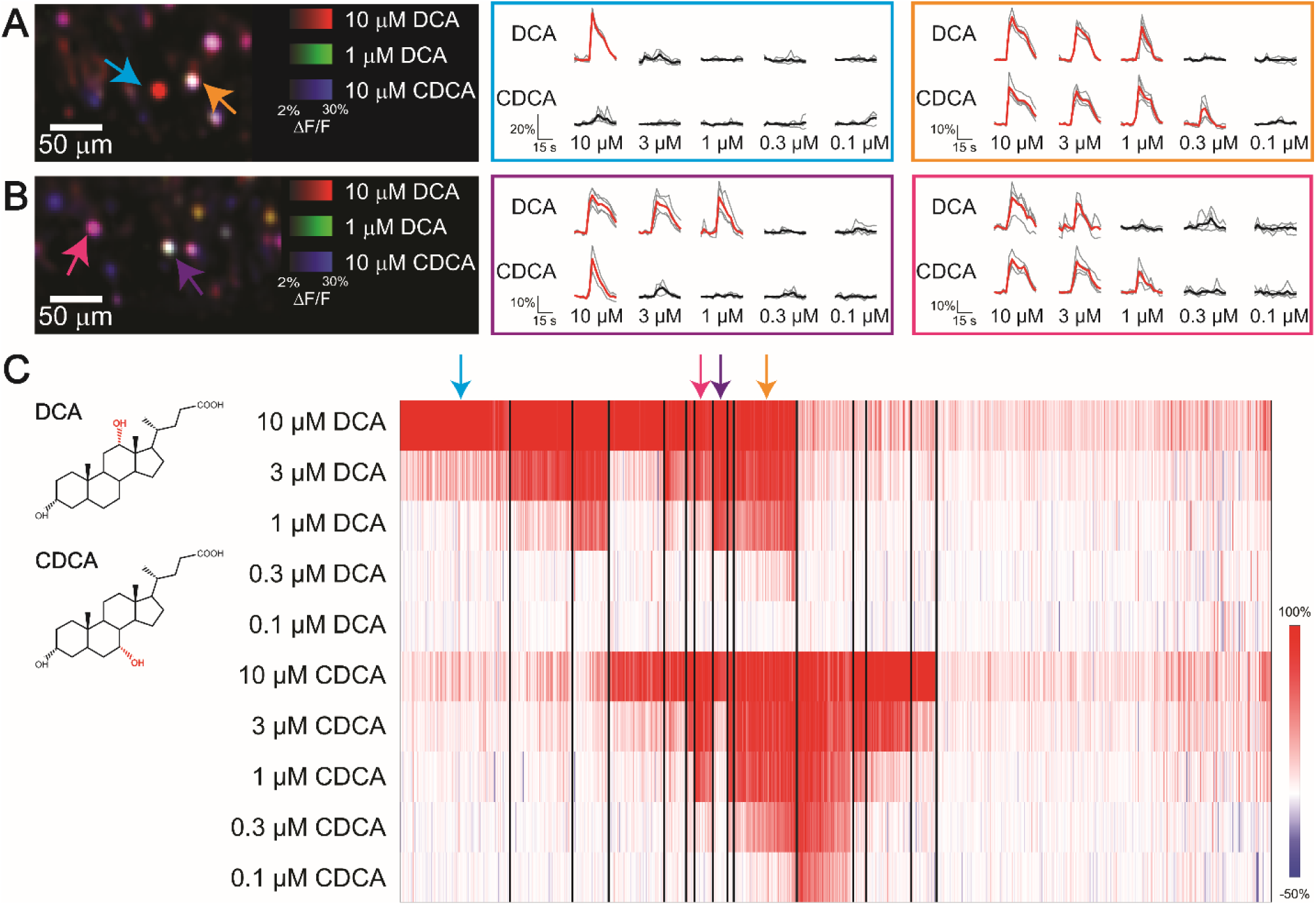
Vomeronasal sensory neuron responses to deoxycholic acid and chenodeoxycholic acid. (**A-B**) Representative colorized images of VSN responses (ΔF/F) in a single frame of an OCPI image stack (left). The responses to 10 µM DCA (red), 1 µM DCA (green), and 10 µM CDCA (blue) are shown in (**A**). The responses of VSNs to 10 µM DCA (red), 1 µM DCA (green), and 10 µM CDCA (blue) are shown in (**B**). Across-trial VSN responses are plotted as individual traces. Bolded trace indicates the mean response across all stimulus repeats. Responses in the cyan, orange, purple, and magenta box corresponds to the VSN indicated by the cyan, orange, purple, and magenta arrow respectively. (**C**) Clustered heat map of VSN response to DCA and CDCA (structures at left) at varying concentrations. Each column indicates an individual neuronal response. Cluster divisions are indicated by black vertical lines. Arrows above the heat map highlight the clusters in which neurons from (**A**) and (**B**) fall. Shown are the responses of 1,018 VSNs from 3 OMPxAi96 animals (2 males and 1 female). All experiments shown have ≥ 3 repeats each.

**Supplementary Figure 3:**
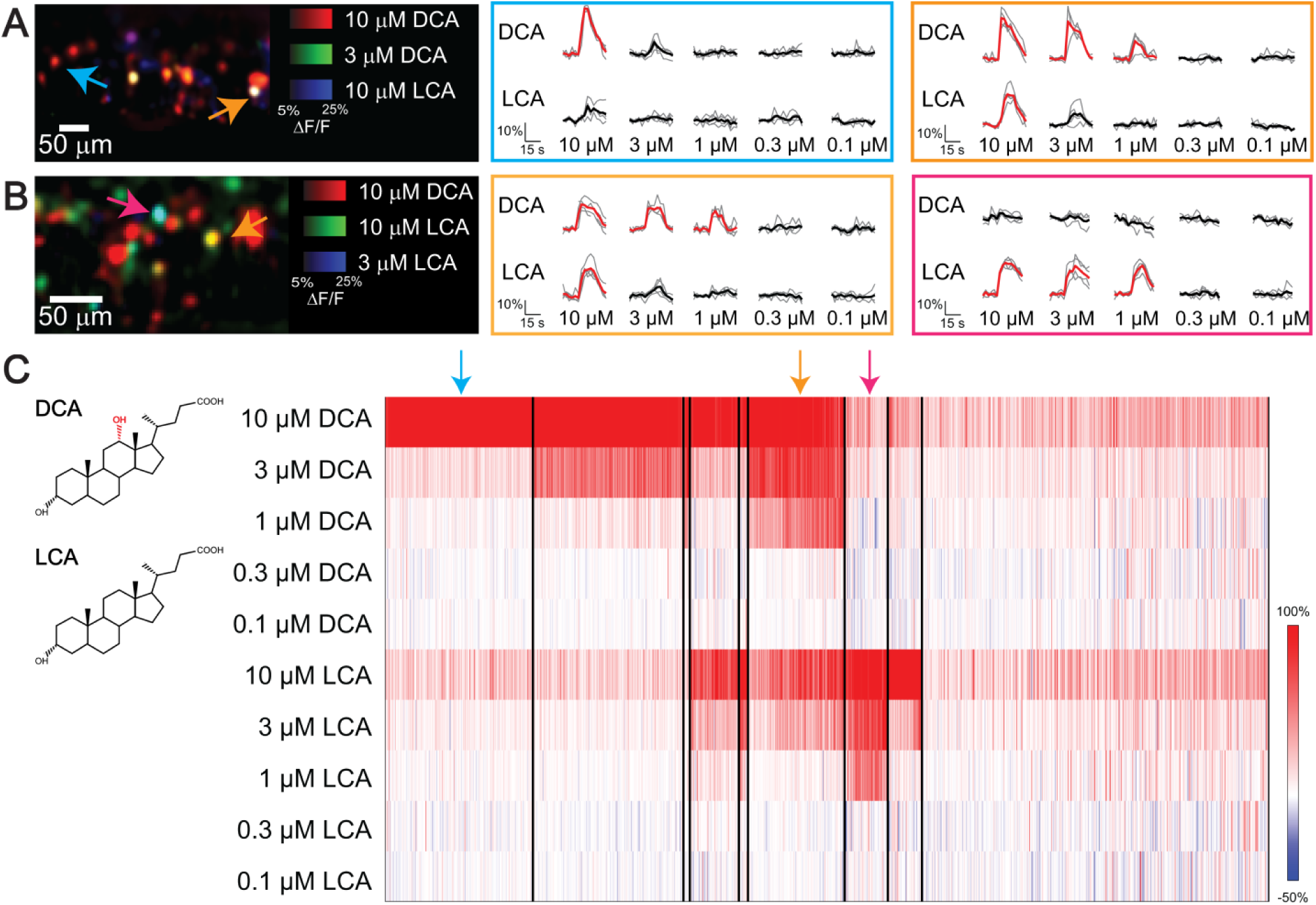
Vomeronasal sensory neuron responses to deoxycholic acid and lithocholic acid. (**A-B**) Representative colorized images of VSN responses (ΔF/F) in a single frame of an OCPI image stack (left). The responses to 10 µM DCA (red), 3 µM DCA (green), and 10 µM LCA (blue) are shown in (**A**). The responses of VSNs to 10 µM DCA (red), 10 µM LCA (green), and 3 µM LCA (blue) are shown in (**B**). Across-trial VSN responses are plotted as individual traces. Bolded trace indicates the mean response across all stimulus repeats. Responses in the cyan, orange, and purple box corresponds to the VSN indicated by the cyan, orange, and purple arrow respectively. (**C**) Clustered heat map of VSN response to DCA and LCA (structures at left) at varying concentrations. Each column indicates an individual neuronal response. Cluster divisions are indicated by thin vertical lines. Arrows above the heat map highlight the clusters in which neurons from (**A**) and (**B**) fall. Shown are the responses of 1,251 VSNs from 3 OMPxAi48 animals (2 males and 1 female). All experiments shown have ≥ 3 repeats each.

**Supplementary Figure 4:**
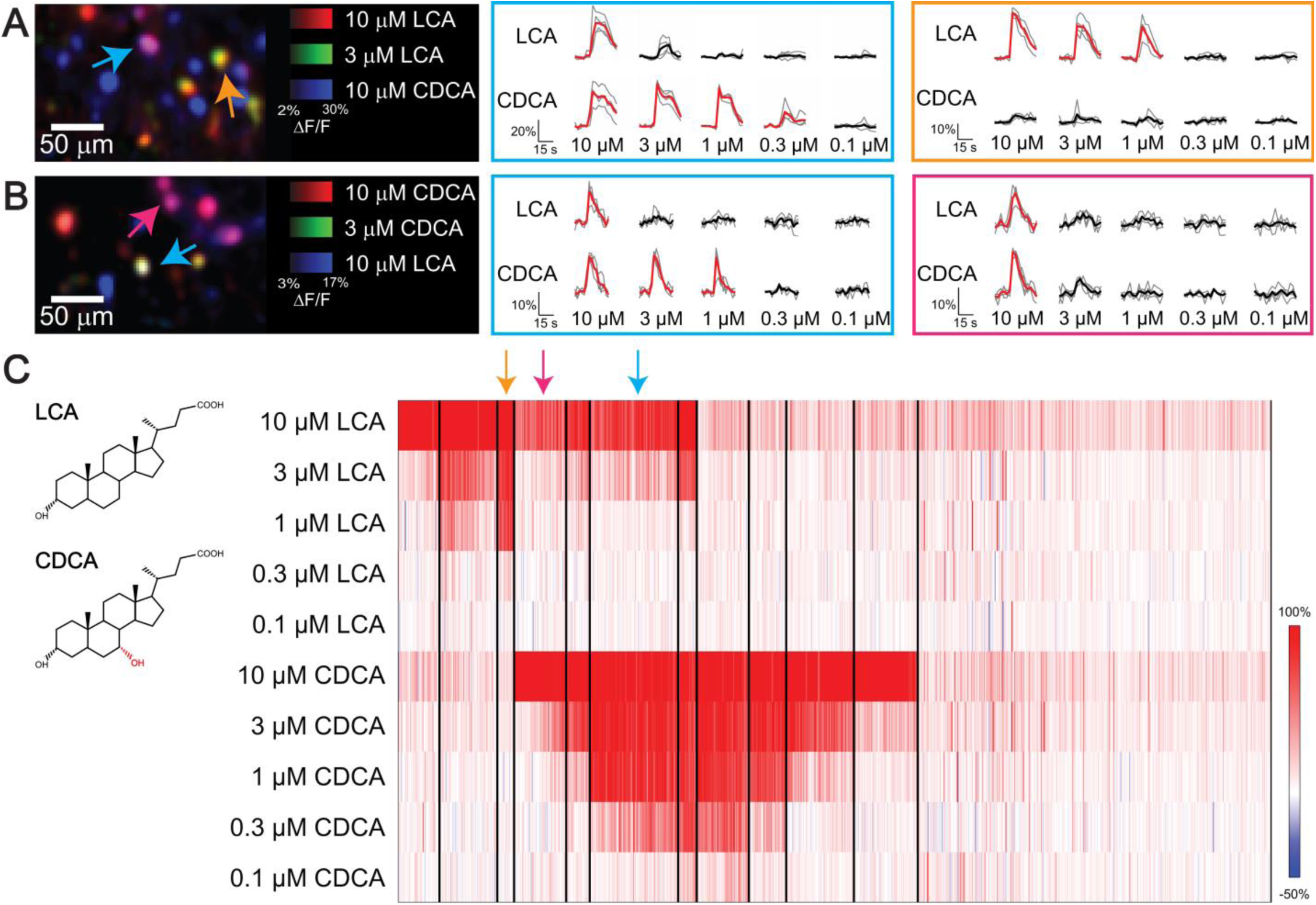
Vomeronasal sensory neuron responses to lithocholic acid and chenodeoxycholic acid. (**A-B**) Representative colorized images of VSN responses to LCA and CDCA (left). (**A**) The responses of VSNs to 10 µM LCA (red), 3 µM LCA (green), and 10 µM CDCA (blue) are depicted. Responses of individual VSNs are plotted as individual traces with the mean response across all stimulus repeats in bold. Responses of the cyan box correspond to the cyan arrow and responses of the orange box correspond to the neuron indicated by the orange arrow. (**B**) Depiction of a single frame where neurons responding to 10 µM CDCA (red), 3 µM CDCA (green) and 10 µM LCA (blue) are shown. Traces in the cyan box or the magenta box corresponds to the neuron indicated by the cyan and magenta arrow, respectively. (**C**) Clustered heat map response of all neurons (each column) imaged in response to varying concentrations of LCA and CDCA (structures at left). Colored arrows over heat map indicates the cluster the representative traces fall under. Shown are the responses of 897 VSNs from 3 animals (2 males, 1 female, OMPxAi148 mice). All experiments shown have ≥ 3 stimulus repeats.

**Supplementary Figure 5:**
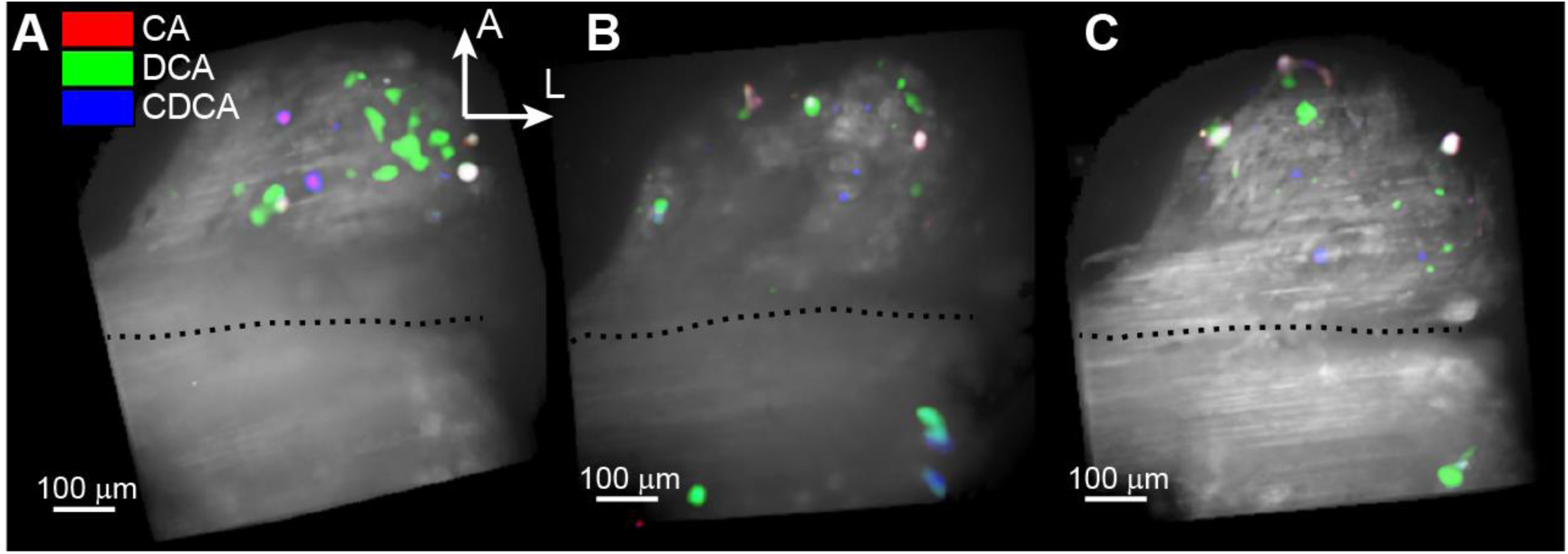
AOB glomerular layer imaging shows distributed BA sensitivity in the anterior (V1R-receiving) subregion. (**A-C**) Maximum projection images of GCaMP6f activation in the AOB glomerular layer of VNO-AOB *ex vivo* preparations. The anterior AOB, which is targeted by V1R-expressing VSNs, is the predominant location where activated presynaptic terminals were reliably found during VNO stimulation with CA, DCA, and CDCA (all at 10 µM). As in VNO imaging experiments, stimuli were delivered in at least 3 randomized interleaved blocks, and colorized regions in these images reflect locations of reliable fluorescence increases.

**Supplementary Figure 6.**
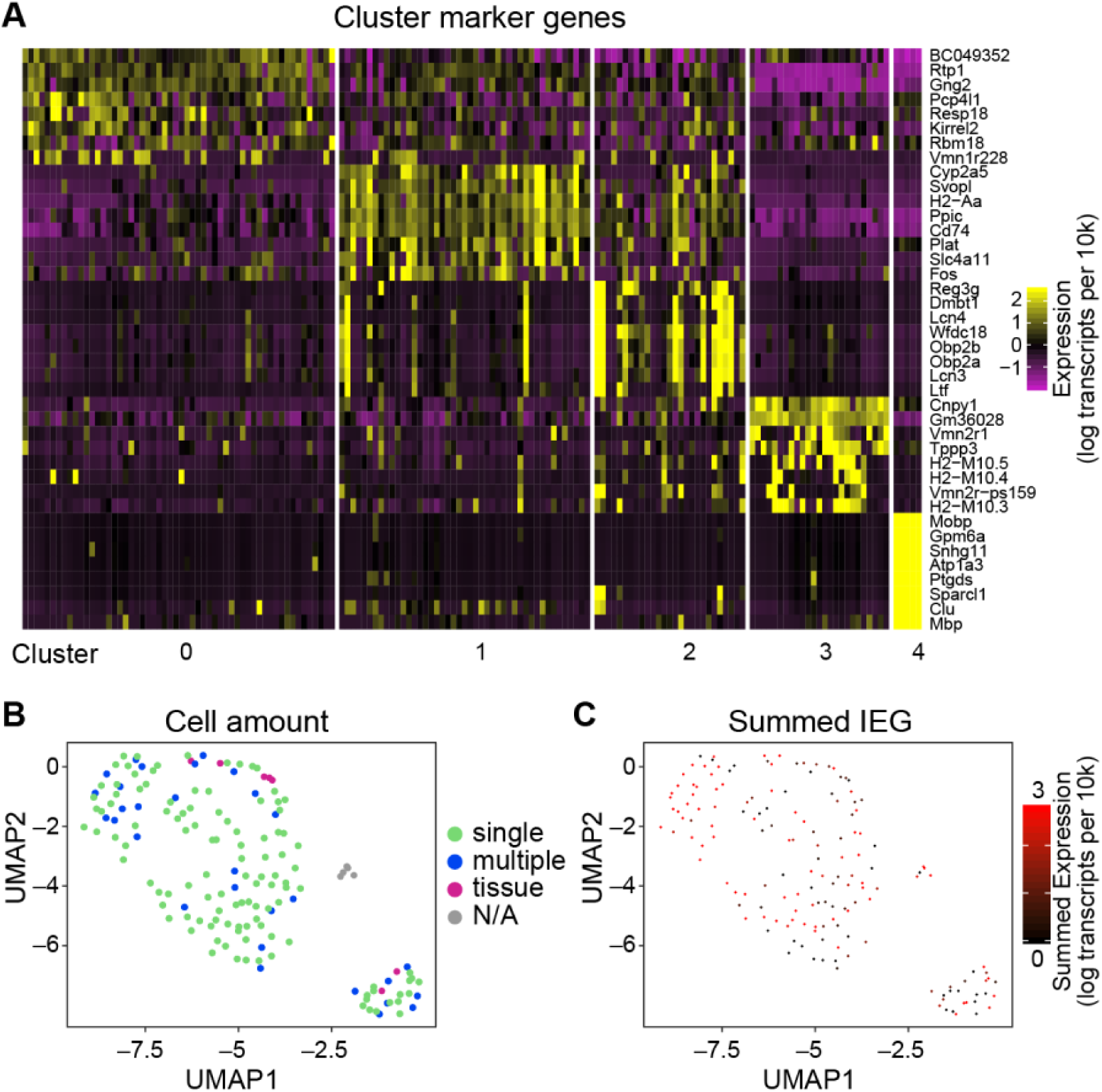
Additional information about VSN single cell RNAseq analysis. (**A**) Top 8 marker genes for each cluster shown in Figure 5. Marker genes were selected based on the Wilcoxon rank sum test (normalized expression, log transcripts per 10k). (**B**) UMAP plot with each sample colorized based on the user-reported number of visible cells (or cell-like material) attached to the patch pipette during the FLICCR-seq procedure. No apparent enrichment for single versus multiple cells was seen in any specific cluster. (**C**) UMAP plot showing the summed immediate early gene expression (log transcripts per 10k) for each sample. Genes included in this IEG pool are *Egr1, Fos, Jun, Arc, Fosb*, and *Nr4a1*. IEG expression could reflect recent activation by stimuli (*i.e*., the bile acids used to identify bile acid-responsive cells) or the VNO slicing procedure (or both).

**Supplementary Movie S1**: Volumetric view of OCPI imaging of GCaMP6s-expressing VNO tissue. The same tissue is shown in 3 orthogonal views and an isometric view. The volumetric stack acquisition rate was 0.3 Hz (1 frame every ∼3 s).

**Supplementary Movie S2**: VNO slice imaging of GCaMP6s-expressing VSNs for FLICCR-seq. Synchronized neuronal activity during slower epochs corresponds to times during which 1 µM CA or 1 µM DCA was applied. Periods during which spontaneous VSN flickering appears rapid are fast-forwarded at ∼10x (inter-stimulus periods).

